# Growth compensation upon changes in tissue size in the *Drosophila* abdomen

**DOI:** 10.1101/2024.10.02.616326

**Authors:** Ana Ferreira, Andrea Cairoli, Federica Mangione, Maxine V. Holder, Anna Ainslie, Birgit L. Aerne, Guillaume Salbreux, Nicolas Tapon

## Abstract

Attaining the appropriate size during development is essential for the function of animal tissues and organs. Robust tissue size control implies the existence of compensatory mechanisms that allow developing systems to recover from growth perturbations. However, the difficulty of directly observing normal or compensatory developmental growth means we have little understanding of the cellular behaviours that confer robustness to tissue size control. Here, we study how growth perturbations affect proliferation kinetics and the timing of growth termination of *Drosophila* histoblasts, the progenitors that give rise to the adult abdominal epidermis. Histoblasts undergo extensive growth and proliferation during the pupal stage, which is accessible for long-term live-imaging and precise quantitative analysis. By manipulating cell number or volume prior to the pupal growth phase, we changed the starting size of the abdomen primordium, then observed how the histoblasts adapted to these changes by altering their growth dynamics. We show that, upon a decrease in starting tissue size, the histoblasts compensate by extending their temporal proliferative window, undergoing additional cell cycles, as well as increasing their apical area to maximise coverage of the abdominal surface. When initial tissue size is increased, the histoblasts undergo fewer division cycles and arrest proliferation earlier than normal. Thus, the proliferative window of this tissue is flexible enough to buffer for changes in tissue size. Our data also suggest that the histoblasts sense both spatial and temporal cues to arrest their growth at the appropriate time and ensure accurate tissue size control.

## INTRODUCTION

Developmental growth control in animals is essential for each tissue and organ to achieve the correct size and shape^1–3^. During development, organs grow until they reach their characteristic final size, at which time most cells stop proliferating except for stem or progenitor cells responsible for maintaining adult homeostasis and wound repair^1–3^. The striking robustness of body and organ size across individuals of the same species suggests that control mechanisms sense tissue size to both terminate developmental growth at the appropriate time and respond to perturbations. One example of this robustness is regenerative growth, whereby some adult and many developing tissues have the capacity to respond to acute cell loss and can regenerate an almost normal structure by formation of a regenerative blastema that undergoes growth and repatterning^4–6^.

Resilience in response to changes in tissue size has been experimentally investigated by manipulating the size of the primordium before its expansion to test whether the tissue is able to adapt. Early mouse embryos can be micromanipulated or disaggregated and reaggregated into morulas of larger or smaller sizes that are then transferred into pseudo-pregnant females and allowed to develop^7–10^. These studies have demonstrated that growth compensation occurs uniformly across the embryo but at different times (E5.5-6.5 for larger embryos and gastrulation or later for smaller embryos), though there are no mechanistic insights into this process. Interestingly, later during development, some organs (e.g. liver, heart) are able to undergo growth compensation in response to changes in primordium size, while others cannot (e.g. pancreas)^11,12^. Compensatory, or catch-up growth has also been observed in human children following periods of growth restriction, suggesting this is a widely conserved phenomenon^13,14^. Importantly, by challenging a developing system and carefully examining the parameters of compensatory growth, it should be possible to gain insights into the size-sensing mechanisms responsible for growth termination under normal conditions. However, as direct long-term observation of developmental growth remains difficult, we have very limited understanding of the cellular behaviours that allow compensatory growth and their relationship to “wild type” tissue size regulation.

The *Drosophila* abdominal epidermis is an ideal system to study these fundamental questions *in vivo*, since it is amenable to live-imaging, and we have previously developed an analysis pipeline that allows a comprehensive quantitative analysis of growth dynamics^15^. The adult abdominal epidermis originates from imaginal cells, specified during embryogenesis, called histoblasts^16^. These cells are arranged in small clusters (nests) embedded in the larval epidermis (Figure 1A). Each abdominal hemisegment contains two nests located dorso-laterally (dorsal anterior and dorsal posterior), one located ventrally, and one spiracular nest located laterally^17,18^. During the larval period, the histoblasts remain quiescent and accumulate mass while arrested in G2^19,20^ (Figure 1A). At the onset of pupariation (0h After Puparium Formation - APF), a pulse of the steroid hormone ecdysone triggers the histoblasts to enter the cell cycle^21^. During this stage (prepupal stage) the histoblasts undergo three rapid synchronous cleavage (growthless) divisions^17^ (Figure 1A). This first stage of cell proliferation occurs between 0-10h APF, with an average cell cycle time of ∼3h^15,22^. At around 14h APF, the histoblasts begin an expansion phase where they undergo cell growth and an average of four cell divisions with a longer doubling time (∼4.5h)^15,22^. During this phase, the histoblasts expand to replace the surrounding larval tissue until the fusion of the right and left side nests at the dorsal midline. The larval epithelial cells (LECs), which formed the larval epidermis, extrude basally from the epithelial layer and undergo apoptosis^21^. At around 24h APF, the histoblasts start to exit the cell cycle and by 30h APF most cells have stopped dividing, therefore growth termination occurs through an abrupt transition to cell cycle arrest^15^. Differentiation then takes place to form the adult abdominal epidermis (Figure 1A).

**Figure 1.**
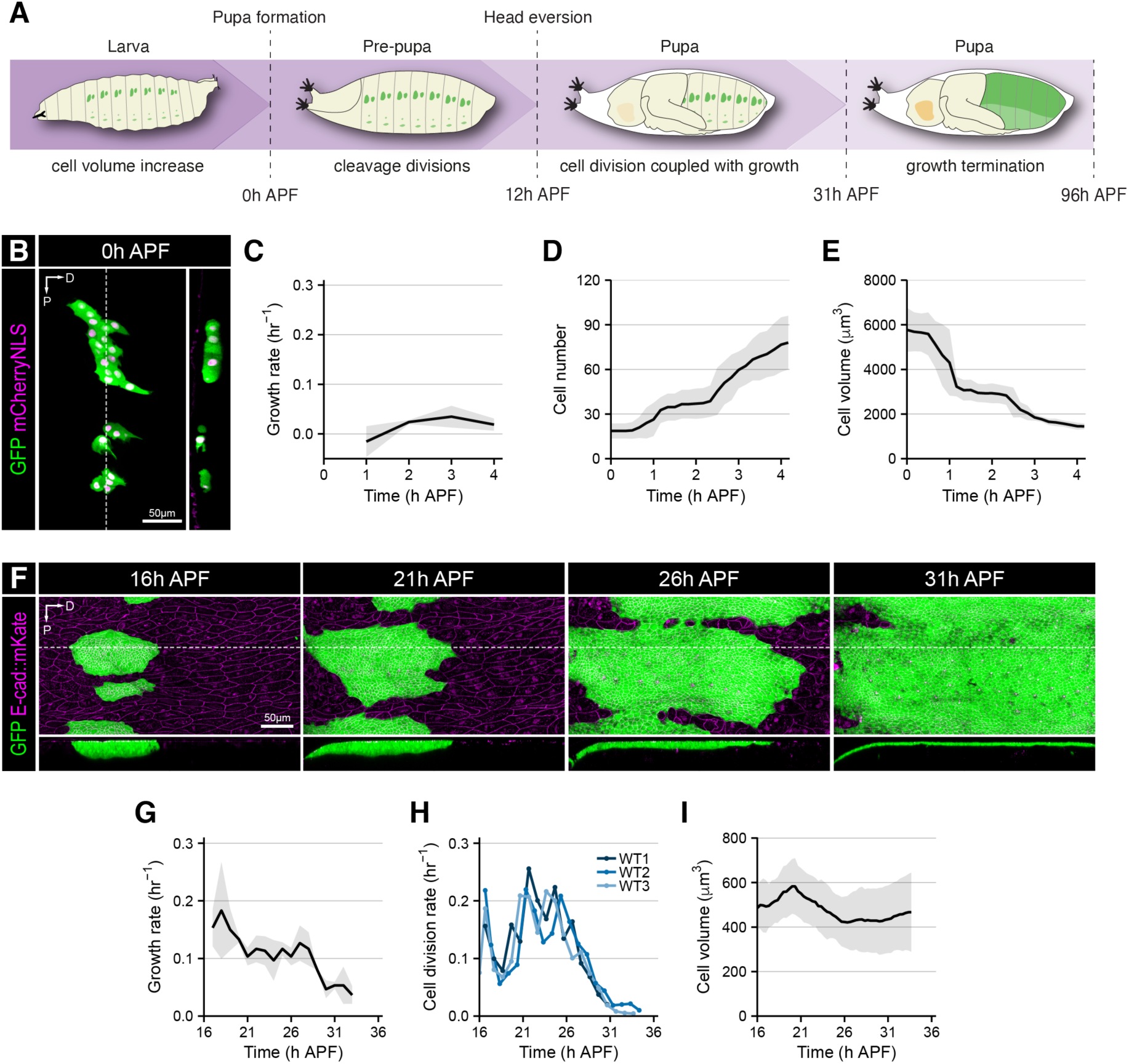
Analysis of abdominal tissue growth dynamics. (A) Schematic of the different phases of abdominal epidermis development. (B) Confocal image of histoblast dorsal nests at 0h APF. Histoblasts were labelled by driving *nls-mCherry* (in magenta) and a ubiquitous *EGFP* (in green) in this tissue. Dashed white line indicates where the cross-section shown on the right side was taken. Note that the posterior nest is sometimes split into smaller islands, which fuse during the cleavage cycles. Scale bar = 50 μm. Dorsal (D) is to the right; anterior is to the top; posterior (P) to the bottom. See also Video S1. (C) Growth rate as a function of time during the first 4h of prepupal development (n=3). Growth rate was calculated as the tissue volume difference between two timepoints divided by initial tissue volume, divided by the time difference between the two time points. (D) Cell number counts during the first 4h of the prepupal stage (n=3). (E) Average cell volume during the first 4h of the prepupal stage. Cell volume was calculated by dividing total tissue volume by the number of cells for each timepoint (n=3). (F) Confocal images of histoblasts labelled by driving expression of a ubiquitous EGFP (in green) in the tissue and expressing *E-cad::mKate* (in magenta) and the time points indicated. White dashed lines indicate the position of the cross sections shown in the panels below. Scale bar = 50 μm. See also Video S1. (G) Growth rate as a function of time (n=3). Growth rate was calculated as noborder ROI tissue size difference between two timepoints divided by initial tissue size. Data from noborder ROIs. See also Figure S1, Video S1 and STAR Methods. (H) Cell division rate in each wild-type movie as a function of time. Data from noborder ROIs. (I) Average cell volume as a function of time (n=3). Cell volume was calculated by dividing the total volume by the number of cells for each timepoint. Shaded grey error bars represent SD.

The target tissue size (total surface the histoblasts must cover in each segment) is determined by the volume of the pupa at the end of the larval growth period. Thus, the fly abdomen is an ideal tissue to study growth compensation: we can alter starting primordium size without changing target tissue size by manipulating the number or size of histoblasts during the larval stages and observe how the cells adapt to this challenge. Using this approach, we show that the timing of proliferation arrest can change upon perturbing initial histoblast nest size: reduced histoblast number leads to delayed arrest, while increased number triggers earlier growth termination. This suggests that the histoblasts can sense tissue size and adjust the timing of growth termination to ensure the full occupancy of the abdomen surface is achieved.

## RESULTS

### Measuring growth dynamics during abdominal epidermis development

We previously created an image analysis pipeline to precisely track cells and quantify their division kinetics during abdominal epidermis growth based on imaging the junctional marker E-cad::GFP^15^. To assess growth, volume measurements are also needed, therefore we established a method to track tissue volume as well as apical surface. We used Imaris to reconstruct the surface of the tissue (STAR Methods; Figure S1A; Video S1). For the early prepupal stage (Figure 1A) we expressed both a ubiquitous GFP and a red nuclear marker specifically in the histoblast nests (Figure 1B). We followed the cleavage divisions until ∼4h APF (Video S1) and observed that, as expected^17^, tissue volume remains constant during this period (Figure S1B). Thus, global tissue growth rate during this period is close to zero (Figure 1C). Cells do undergo cleavage divisions (Figure 1D), leading to decreased cell size, with cells halving their size with each round of division (Figure 1E), which is reflected by a negative cell growth rate during this period (Figure S1C).

Next, we measured growth rates during the expansion phase, when the histoblasts divide and grow to cover the abdominal surface. We imaged both cell junctions with *E-cad::mKate2* and a ubiquitous GFP expressed specifically in the histoblasts for volume reconstruction from ∼16h APF until ∼34h APF (STAR Methods; Figure 1F; Video S1). The emergence of the sensory organ precursor cells (SOPs) was used to temporally align our movies (STAR Methods; Figure S1D-F). During this period, the histoblasts transition from a pseudostratified columnar organisation to an unstratified cuboidal shape (Figure 1F). We defined an anterior “no border region of interest” (noborder ROI) comprising complete lineages of anterior nest histoblasts that never contact the edge of the image frame. To calculate growth rates, we measured the volume of the noborder ROI throughout development (Figures S1G). Tissue growth rates during this period are initially high, then they level out for most of the duration of the expansion phase until they start to decrease at ∼28h APF (Figures 1G, and S1H). The drop in tissue growth rate coincides with the time at which proliferation starts to decrease and cells transition to an arrested state^15^ (Figures 1G, H and S1I). At the cellular level, we observed that average cell volume remains largely constant (Figures 1I and S1J) compared with the cleavage phase (Figure 1E). Thus, in wild type animals, tissue growth starts at the transition between the cleavage and expansion divisions, before decaying as the cells enter cell cycle arrest.

TOR inhibition reduces initial tissue size and leads to a delay in proliferation arrest To address the ability of histoblast nests to compensate for alterations in initial tissue size, we devised experimental conditions that change histoblast number or size. We first reduced cell size by blocking TOR (target of rapamycin) signalling in the histoblasts. The TOR pathway is a conserved signalling cascade that drives cellular growth from yeast to mammals^23,24^. Overexpression of the negative regulators TSC1 and TSC2 (Tuberous Sclerosis Complex 1 and 2) in flies is sufficient to reduce cell growth by inhibiting the mTORC1 complex (mechanistic Target of Rapamycin Complex 1)^25–28^. We blocked TOR signalling in the histoblasts by driving the expression of *UAS-TSC1, UAS-TSC2* transgenes^25^ from the embryonic stage using the *esg^NP^*^1248^–*GAL4* driver. While TOR signalling is active in control animals from 0h APF until 31h APF, as shown by the lack of expression of the *unk* reporter^29^ (Figure S2A), TSC1 and TSC2 expression in the histoblasts was able to inhibit TOR activity from 0h APF until ∼21h APF (Figure S2B). From ∼21h APF the activity of *esg^NP^*^1248^– *GAL4* driver decreases and TOR inhibition is relieved (Figure S2B).

TOR inhibition in the histoblasts led to a reduction in tissue volume at 0h APF (Figures 1B, 2A, and 2B) due to reduced cell size, while cell number was comparable to controls (Figure 2C-E). Thus, TOR signalling is not necessary for histoblast specification during embryogenesis but is required during the larval stages for histoblast cell growth. Tissue size upon TOR inhibition, although reduced, remained constant during the prepupal stage of development (Figure 2B) with tissue growth rates close to zero as in wild-type animals (Figure 2F). However, TOR-inhibited cells largely failed to undergo cleavage divisions, maintaining a more stable volume than controls (Figure 2C-E, and S2C). Cell number therefore increased little in the first four hours of pupal development in contrast to controls where histoblasts divide twice during the same time span (Figure 2C-E, and Video S2). Thus, TOR inhibition reduces cell volume growth during the larval stages and leads to a decrease in the number of early cleavage divisions during the first hours of pupal development.

**Figure 2.**
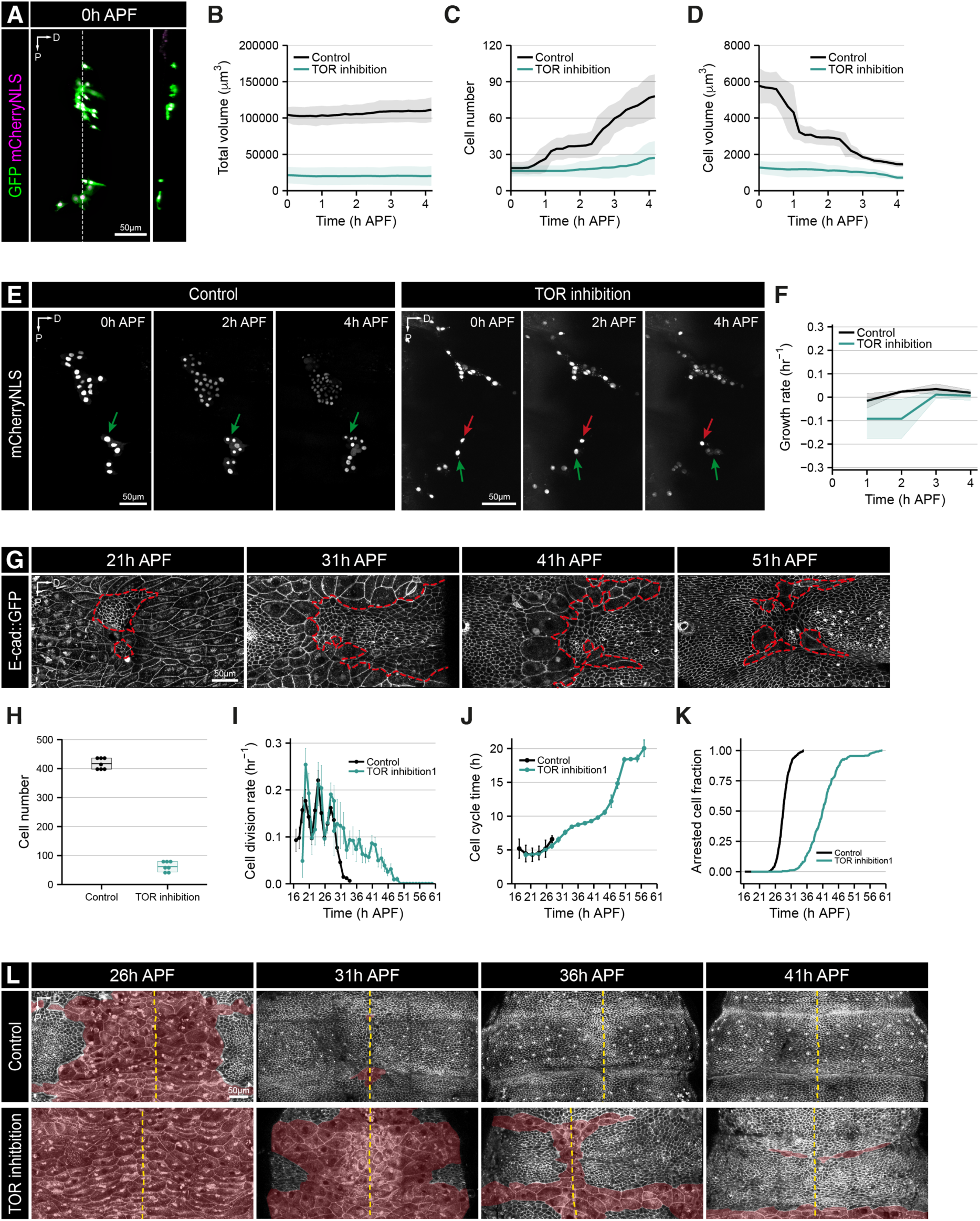
Effects of TOR inhibition on histoblast growth dynamics. (A) Confocal image of histoblast dorsal nests at 0h APF upon inhibition of TOR signalling. Histoblasts were labelled by driving *nls-mCherry* (in magenta) and a ubiquitous *EGFP* (in green) in this tissue. Dashed white line indicates where the cross-section shown on the right side was taken. Scale bar = 50 μm. Dorsal is to the right; anterior is to the top. See also Video S1. (B) Total tissue volume as a function of time in wild-type histoblasts and upon inhibition of TOR signalling (n=3). Error bars represent SD. (C) Quantification of cell number during the first 4h of the prepupal stage (n=3). Error bars represent SD. (D) Average cell volume during the first 4h of the prepupal stage. Cell volume was calculated by dividing total tissue volume by the number of cells for each timepoint (n=3). Error bars represent SD. (E) Example time-lapse confocal images of histoblasts expressing *nls-mCherry* at the time points indicated. Left side panels are control animals while right side panels represent pupae where TSC1/TSC2 were overexpressed in the histoblasts. Green arrows highlight divisions, while red arrows show cells that did not undergo any divisions during the imaging period. Scale bar = 50 μm. Dorsal is to the right; anterior is to the top. See also Video S1. (F) Growth rate as a function of time during the first 4h of prepupal development (n=3). Growth rate was calculated as the tissue size difference between two timepoints divided by initial tissue size. Error bars represent SD. (G) Time-lapse confocal images of a pupa expressing *E-cad::GFP* and overexpressing TSC1/TSC2 in the histoblasts at the time points indicated. Histoblasts are outlined in red. Scale bar = 50 μm. Dorsal is to the right; anterior is to the top. See also Video S1. (H) Cell number counts at 16h APF for wild-type and TOR inhibition movies (n=7). (I) Cell division rate in a wild-type and a TOR inhibition movie as a function of time. Data from noborder ROIs. Error bars represent the SEM. See also Figure S1 and Methods S1. (J) Cell cycle time as a function of developmental time for wild-type and TOR inhibition movies. The dots represent the binned data, and the error bars are the SEM for the bin. Data from noborder ROIs. (K) Fraction of arrested cells calculated as the time of final division, as a function of time, for wild-type and TOR inhibition movies. Data from noborder ROIs. (L) Time-lapse confocal images from a dorsal view of pupa expressing *E-cad::GFP* in a wild-type background (left panels) or upon TOR inhibition (right panels) at the time points indicated. LECs are shadowed in red and the dashed yellow line indicates the position of the dorsal midline. Scale bar = 50 μm. Dorsal is to the right; anterior is to the top.

Is TOR activity directly required for the cleavage divisions during the prepupal stage or do the cleavage divisions fail to occur because of reduced cell growth during the larval stages? To address this question, we performed temporally controlled TOR inhibition experiments with *tub-GAL80^ts^* ^30^ and used the adult abdominal phenotype as a readout. When TOR is inhibited continuously with the *esg^NP^*^1248^–*GAL4* (from embryonic stages until ∼ 21h APF; Figure S2B), adult flies have a normal abdomen in terms of histoblast coverage (as visualised by the normal pattern of cuticle deposition), but the number of SOPs is reduced and spacing between them is uneven (Figures S2D and S2E). Confining TOR inhibition from the embryonic stage to the end of the larval period results in similar adult abdominal defects (Figure S2F). In contrast, inhibition of TOR either from the end of larval development or during the expansion phase (from ∼8-12h APF) results in a normal adult abdomen with no defects in either SOP number or pattern, although animals could not hatch, presumably because of TOR inhibition elsewhere in the animal (Figures S2G and S2H). These results suggest that the consequences of TOR inhibition on histoblast proliferation during the pupal stages are due to the absence of cell growth during larval development. Histoblasts need to accumulate Cyclin E (CycE) during the larval stages to be able to undergo the three fast cleavage divisions upon pupariation^22^. TOR signalling has been shown to affect CycE levels in other *Drosophila* tissues^25^. Inhibition of TOR in histoblasts might therefore prevent CycE accumulation during the larval period, thereby preventing rapid cleavage during the prepupal stage. In addition, TOR inhibition may impair cleavage by reducing cell volume to an extent where division is no longer possible.

We then assessed the consequences of TOR inhibition on histoblast expansion. As for wild-type animals (Figure 1 and ^15^), we analysed the expansion phase upon inhibition of TOR signalling using long-term live imaging of cell junctions labelled with *E-cad::GFP*, followed by cell segmentation and tracking (Figure 2G; Video S2). Movies of the TOR-inhibited condition were temporally aligned to controls based on SOP emergence (^15^, STAR Methods and Figure S3A, S3B). As a result of the reduction in cleavage divisions, expansion in TOR-inhibited animals starts with a lower cell number (∼50 versus ∼400 in controls at 16h APF, Figure 2H). We first examined the cell division rate (number of cell divisions per unit of time, relative to the total number of cells) from 16h APF onwards. In wild-type animals, the cell division rate shows an oscillatory behaviour with a ∼4h period, with three peaks observed before starting to decrease at ∼28h APF, and cell divisions stopping by ∼32h APF (Figures 1H, 2I, 2J and ^15^). Upon TOR inhibition, while we still observed three oscillatory peaks in cell division rate, the final decrease in cell division rate was more gradual, with cell divisions persisting ∼15h later than in control animals (Figures 2G, 2I and S3C). Strikingly, while the cell cycle time (time between two divisions) is similar to wild-type cells until 28h APF, it increases up to 20h during the extended proliferation window (Figure 2J and Figure S3D).

Is the timing of cell cycle exit affected by TOR inhibition? We computationally labelled cells as they performed their final cell division (arrested cells)^15^. In wild-type animals, half of the cells are arrested at ∼25h APF and after 28h APF nearly all newly created cells are arrested^15^ (Figures S1I and 2K). In the TOR-inhibited condition, cell cycle exit is considerably delayed, with half of the cells arrested at ∼39h APF, and the transition to growth termination is slightly slower, as dividing cells persist until ∼48h APF (Figures 2K and S3E-G).

To test whether the delay in cell cycle arrest correlates with a delay in covering the entire abdominal surface, we imaged the dorsal surface of the abdomen, since the replacement of the dorsal LECs occurs at the final stages of histoblast expansion (Figure 2K). In wild type animals, fusion of the histoblast nests at the dorsal midline occurs around 30-32h APF, consistent with previous work^31^. In TOR-inhibited animals, LEC replacement by the histoblasts was markedly delayed and continued beyond the normal time window until the histoblasts met at the midline, at ∼41h APF (Figure 2L). This is consistent with the idea that histoblasts respond to spatial cues, delaying cell cycle exit when the abdominal segment surface has not been covered. Furthermore, we used laser ablations at 0h APF to confirm previous cauterisation experiments^18^ showing that histoblasts do not cross the dorsal midline or intersegmental boundaries (Figure S3H). This provides further evidence that histoblasts are able to sense and respect the spatial boundaries of their segment.

How is the number of divisions affected by TOR inhibition? Wild-type histoblasts undergo an average of 7 divisions during pupal development: three cleavage divisions and four expansion divisions^15^. Only three of the expansion divisions can be live-imaged (Figures 3A and 3B), as the last cleavage and the first expansion division occur during large-scale morphogenetic movements including head eversion, which preclude continuous imaging between 4 and 16h APF. Upon TOR inhibition, the number of divisions before 16h APF was more than halved (∼1.56 +/-0.5 compared with ∼4.05 +/-0.3 divisions for controls during the same time period). However, we were able to observe more divisions during the expansion phase from 16h APF (∼4.8 +/-0.8 divisions compared to ∼2.9 +/- 0.55 in controls) (Figures 3A-D and S3I). Thus, TOR inhibition limits the number of cleavage divisions by reducing cell growth during the larval stages, but the histoblasts can compensate by delaying cell cycle exit and undergoing additional expansion divisions.

**Figure 3.**
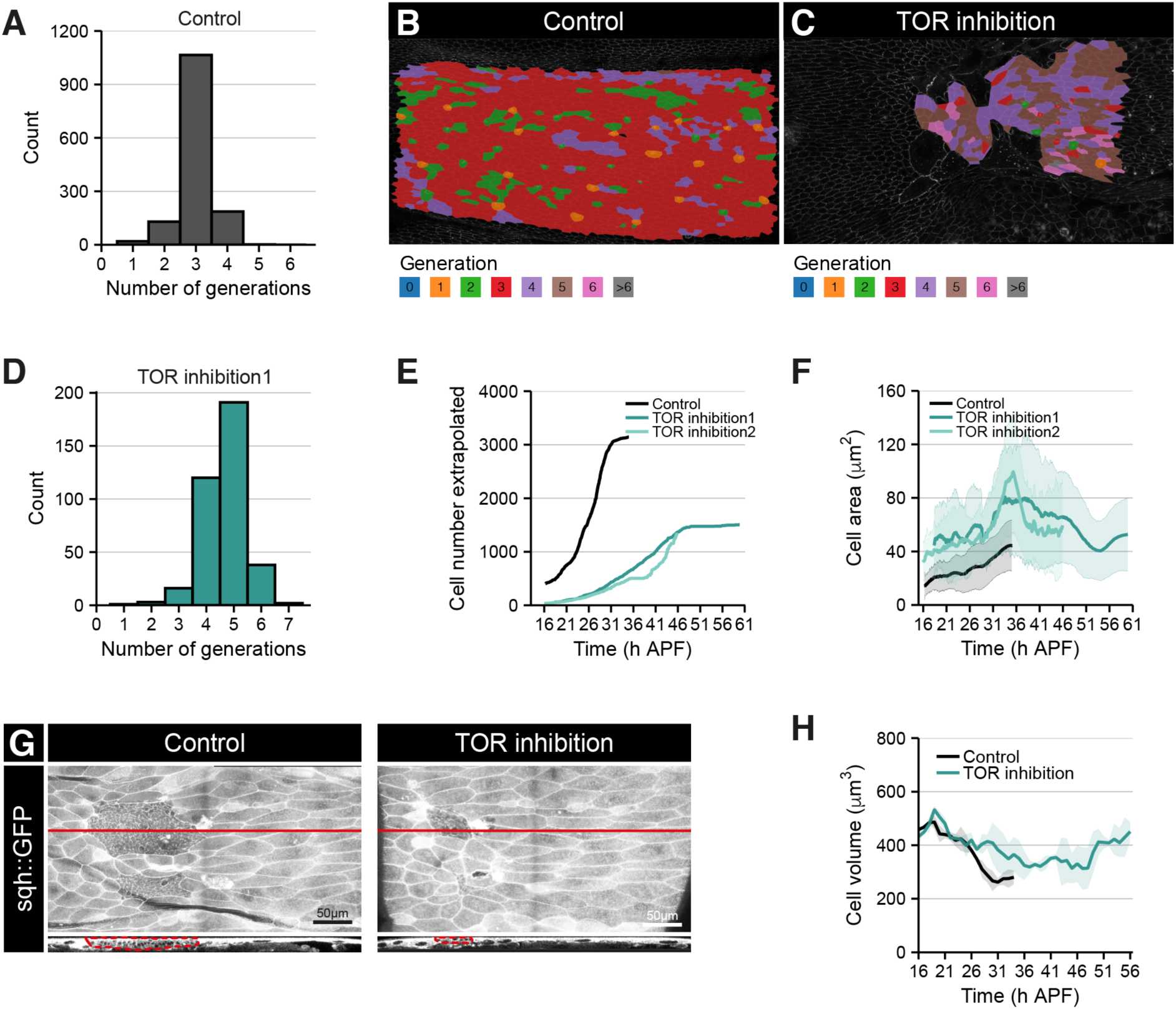
TOR inhibition causes cells to proliferate more during expansion phase. (A) Histogram showing the distribution of histoblast generation numbers during the expansion imaging window for a wild-type movie. Data from noborder ROI. (B and C) Snapshots of the final frame for a wild-type movie (B) and a TOR inhibition movie (C), with cells coloured according to the number of generations in the noborder ROI. Dorsal is to the right; anterior is to the top. (D) Histogram showing the distribution of histoblast generation numbers during the expansion imaging window for a TOR inhibition movie. (E) Extrapolated cell numbers for the anterior nest as a function of time, for wild-type and TOR inhibition movies. See Methods S1. (F) Apical cell area as a function of time for wild-type and TOR inhibition movies. Data from noborder ROIs. Error bars represent SD. (G) Example confocal images of histoblasts expressing *sqh::GFP* in control animals (left panel) or in animals expressing TSC1/TSC2 in the histoblasts (right panel). The red horizontal line indicates the position of the cross-sections shown below. Scale bar = 50 μm. Dorsal is to the right; anterior is to the top. (H) Average cell volume as a function of time, for control and TOR inhibition movies. See STAR Methods. Error bars represent SD, n=2.

Is the delay in cell cycle exit sufficient to restore normal cell numbers in TOR-inhibited animals? To estimate the number of histoblasts in the anterior dorsal nest throughout the expansion phase, we extrapolated from cell numbers at 16h APF using the cell division rate information in wild-type and TOR-inhibited animals (Figure 3E). This showed that, due to the much lower starting cell number and despite the extra rounds of expansion divisions, TOR-inhibited dorsal anterior nests only reached ∼1400 cells, compared to ∼3,000 for the control (Figure 3E). As final abdominal size (estimated by measuring pupal volume) is similar to controls (Figure S3J), this suggests that the final abdominal epidermis is composed of fewer cells in TOR-inhibited than in wild-type animals. We observed an increase in average apical cell area in the TOR inhibition movies (Figure 3F) suggesting that cells can cover a greater surface by increasing their apical surface. Finally, we also observed that cells from the ventral nest appear to be pulled into the imaging frame, also contributing to ensuring almost complete coverage of the abdominal surface by the histoblasts (Figure 2G; Video S2).

The increased apical area suggests that the histoblasts thin out in order to cover the necessary surface area in the TOR-inhibited conditions, prompting us to measure cell volume. Due to the decline in *esg^NP^*^1248^–*GAL4* expression after 21h APF, we could not measure volume by driving a cytoplasmic GFP in the TOR-inhibited animals as we had done in Figure 1. We therefore used a *sqh::GFP* transgene to measure tissue height and calculated cell volume in the TOR inhibition condition and matching controls (Figures 3G, S3K, S3L, and STAR methods). Cell volume was comparable to that of wild-type cells (Figure 3H). This suggests the histoblasts can indeed adopt a flatter morphology to compensate for reduced cell number. In summary, histoblasts can compensate for reduced initial primordium size by extending the length of proliferation temporal window and increasing cell area to maximise abdominal area coverage.

### Reducing initial histoblast number delays the time of proliferation arrest

To independently test the idea that a reduction in histoblast nest size at the beginning of the expansion phase leads to a compensatory proliferative response, we used laser ablation to kill histoblasts at 0h APF (STAR methods). We eliminated both dorsal nests except one anterior nest cell (Figure S4A). We also ablated all histoblasts from the neighbouring segments on both sides to prevent these cells from invading the analysed segment (Figure S4B). We made sure the animals survived this treatment and the recovered adult animals showed a melanotic scar at the laser ablation site (Figure S4C).

Consistent with the characteristic three rounds of cleavage divisions during the initial hours of pupal development, we observed that the remaining nest was composed of 9 cells at 16h APF and the area occupied was strongly reduced when compared to wild-type animals (Figure 4A; Video S3). The proliferation dynamics were comparable to those observed in the TOR experiment. Proliferation rates oscillated, and cell cycle times were initially similar to wild-type animals (Figures 4B and 4C). At around 30h APF, when proliferation in controls decline, the oscillations became less pronounced and cell cycle time increased up to ∼9h (Figures 4B and 4C). Cell division rates started to decrease at ∼46h APF (18h later than in wild-type animals) with the last divisions taking place at ∼52h APF (Figures 4B and 4D). As in the TOR experiment, the transition to cell cycle exit was considerably delayed and was more gradual (Figures 4C, 4D and S4D-F). Individual cells underwent more divisions from 16h APF (∼7.2 +/-1.3 divisions compared to ∼2.9 +/- 0.55 in controls) (Figures 4E and 4F), while a similar number of divisions occurred prior to the onset of the expansion phase (∼3.2 divisions). Thus, cells compensate for reduced histoblast numbers by undergoing more divisions during the expansion phase. As in the TOR inhibition condition, anterior nest cell number extrapolation showed that the final cell number was reduced but apical cell area was increased when compared to wild-type animals (Figures 4G and 4H).

**Figure 4.**
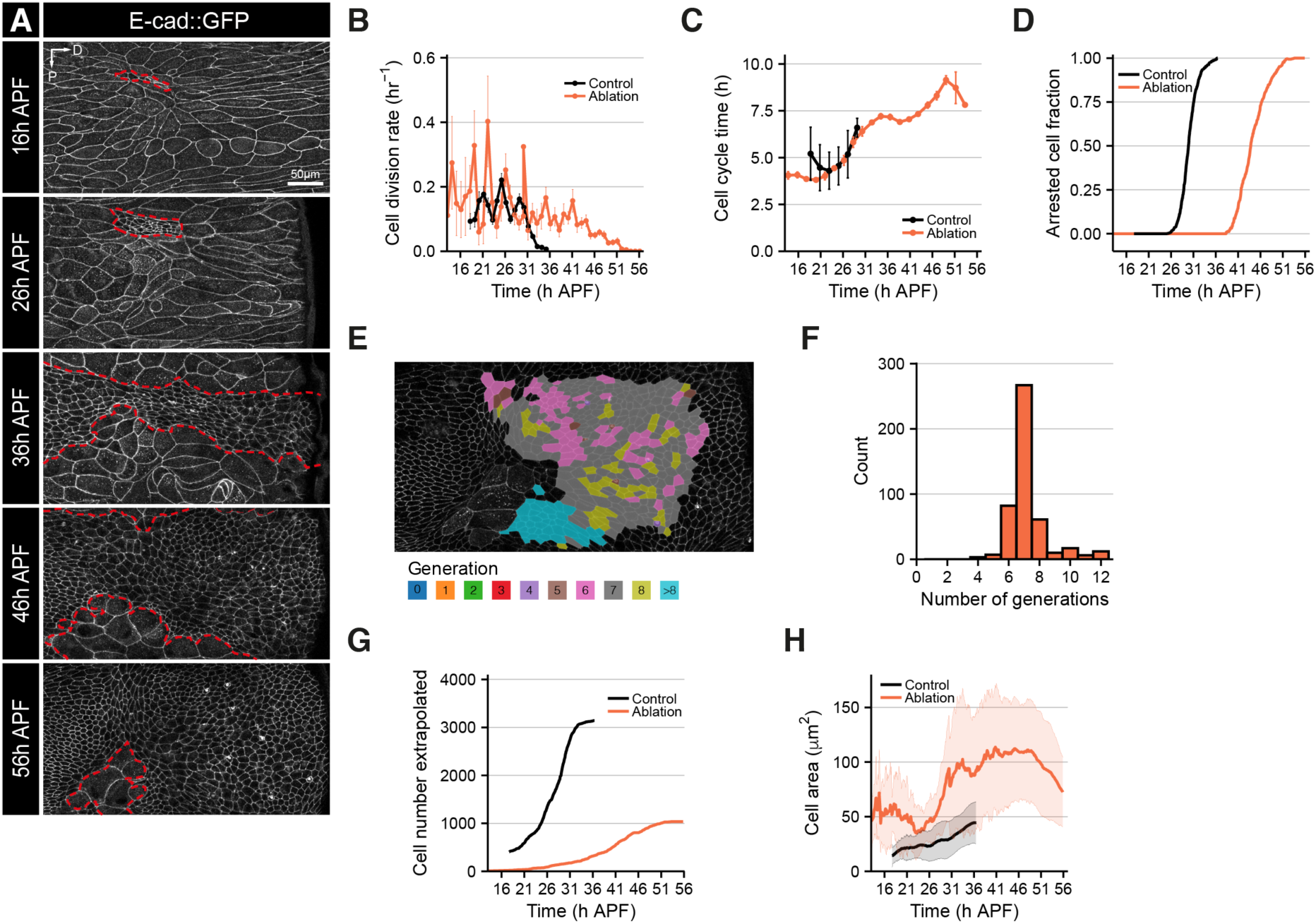
Effects of reducing initial cell number on histoblasts development. (A) Time-lapse confocal images of a pupa expressing *E-cad::GFP* at the time points indicated following laser-ablation of most cells. Histoblasts are outlined in red. Scale bar = 50 μm. Dorsal is to the right; anterior is to the top. See also Video S3. (B) Cell division rate in wild-type and ablation movies as a function of time. Data from the noborder ROIs. Error bars represent SEM. See also Figure S4 and Methods S1. (C) Cell cycle time as a function of developmental time for wild-type and ablation movies. The dots represent the binned data, and the error bars are the SEM for the bin. Data from noborder ROIs. (D) Fraction of arrested cells as a function of time, for wild-type and ablation movies. Data from noborder ROIs. (E) Snapshot of the final frame for the ablation movie, with cells coloured according to the number of generations in the noborder ROI. Dorsal is to the right; anterior is to the top. (F) Histogram showing the distribution of histoblast generation numbers during the expansion imaging window for a wild-type movie. (G) Extrapolated cell numbers for the anterior nest as a function of time, for wild-type and ablation movies. See Methods S1. (H) Apical cell area as a function of time for both wild-type and ablation movies. Data from noborder ROIs. Error bars represent SD.

Together, our results show that starting the expansion phase with a smaller tissue, both upon ablation or by reducing cell size during larval development, leads to a delayed transition to cell cycle arrest and an increase in apical area.

### Increasing histoblast nest size causes premature proliferation arrest

Having shown that reduced starting tissue size could delay cell cycle exit, we wished to test if increasing tissue size could have the opposite effect. Thus, we created larger tissues in an otherwise normal-sized segment by driving the expression of transgenes predicted to increase either cell number or cell size during the larval stages. We used the Gal80^ts^ system^32^ to overexpress either String (Stg)/Cdc25 (a phosphatase that promotes G2/M cell cycle progression^33^), Yki^3SA^ (a constitutively active form of the pro-growth transcriptional co-activator Yorkie^34^), or both, in the histoblasts during the larval stages (STAR Methods; Figures 5A and 5B). All the conditions tested gave rise to significantly larger tissues at the onset of metamorphosis (0h APF) and had little or no effect on animal size (Figures 5B, 5C, and S5A). While Stg overexpression increased cell number but reduced cell size, Yki^3SA^ increased cell size without affecting cell number (Figures 5D and 5E). Combining both Stg and Yki^3SA^ expression increased cell number by a factor of ten-fold, while cell size, though decreased, was closer to the wild-type cell size than Stg overexpression alone (Figures 5D and 5E).

**Figure 5.**
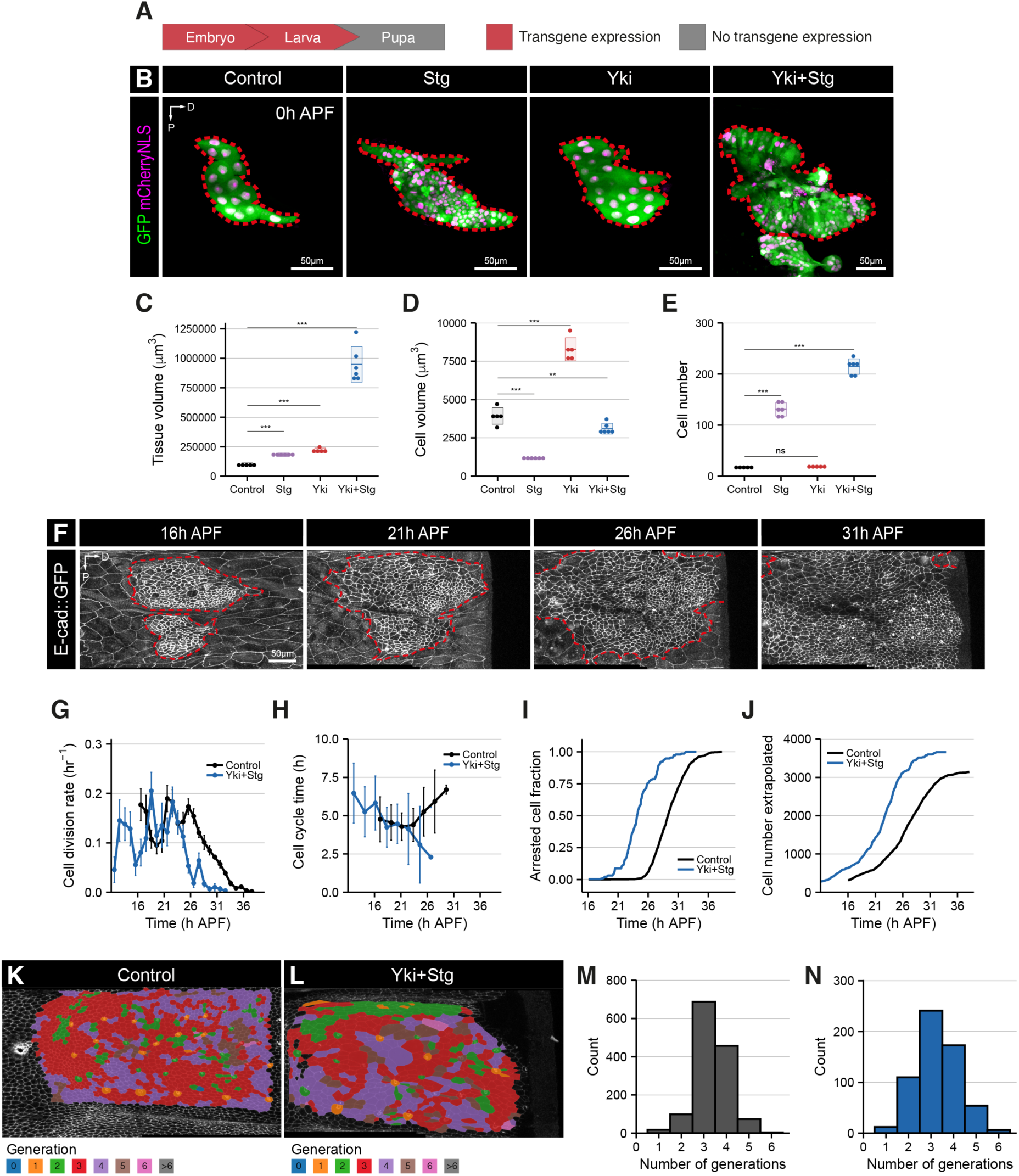
Effects of increasing initial tissue size on histoblast growth dynamics. (A) Schematic depicting the time of transgene expression (in red) or no transgene expression (in grey). (B) Confocal image of histoblast anterior dorsal nests at 0h APF. Histoblasts are labelled by driving *nls-mCherry* (in magenta), a ubiquitous *EGFP* (in green), and either HA (control), Stg, Yki or Yki+Stg in this tissue. Dashed red lines outline the anterior nest. Scale bar = 50 μm. Dorsal is to the right; anterior is to the top. (C) Quantification of tissue volume at 0h APF on all conditions shown in A. (D) Average cell volume at 0h APF for all conditions shown in A. Cell volume was calculated by dividing the total tissue volume by the number of cells. (E) Anterior histoblast nest cell number counts at 0h APF for all conditions shown in A. (F) Time-lapse confocal images of a pupa expressing *E-cad::GFP* at the time points indicated upon expression of Yki+Stg during the larval stages. Histoblasts are outlined in red. Scale bar = 50 μm. Dorsal is to the right; anterior is to the top. See also Video S4. (G) Cell division rate in wild-type and the Yki+Stg overexpression movies as a function of developmental time. Data from noborder ROIs. Error bars represent SEM. See also Figure S4 and Methods S1. (H) Cell cycle time as a function of developmental time for wild-type and Yki+Stg overexpression movies. The dots represent the binned data, and the error bars are the SEM for the bin. Data from noborder ROIs. (I) Fraction of cells that are arrested as a function of time, for wild-type and Yki+Stg overexpression movies. Data from noborder ROIs. (J) Extrapolated cell numbers for the anterior nest as a function of time, for wild-type and Yki+Stg overexpression movies. See Methods S1. (K and L) Snapshots of the final frame for a control movie (K) and Yki+Stg overexpression movie (L), with cells coloured according to the number of generations. Dorsal is to the right; anterior is to the top. (M and N) Histogram showing the distribution of histoblast generation numbers during the expansion imaging window for a control (M) and Yki+Stg overexpression movie (N).

To decide which of these conditions to analyse further, we examined cell numbers and tissue area at 16h APF and found only minor changes upon driving Yki^3SA^ or Stg expression alone (Figures S5B-D). Moreover, upon driving Yki^3SA^ expression, the cell division rate oscillated, with a similar cell cycle time and transition to arrest as control (Figures S5E-I). This suggests that cells compensate during late cleavage/early expansion to normalise tissue size and cell number, and we did not analyse these conditions further. In contrast, co-expression of both Yki^3SA^ and Stg (Yki+Stg) elicited more pronounced phenotypes (Figure 5F; Video S4). Cell division rate oscillated, with an initial cell cycle time longer than in wild-type movies, which progressively decreased to faster cycles than control (Figures 5G-H). However, we note that the final two temporal bins in Figure 5H where cells divided faster contain very few cells (9 and 2) as most had arrested by that time. The decay in the cell division rate occurred several hours earlier than control (Figures 5G and 5I). Accordingly, cell cycle exit occurred earlier, as half of the cells were arrested at ∼24.5h APF, compared with ∼29h APF for wild-type cells (Figure 5I, S5J-L). In this condition, histoblasts divided fewer times than wild-type cells. Comparing cell number at 0h APF (215 cells; Figure 5E) and 14h APF, which marks the beginning of our expansion movie, (282 cells; Figures 5F and 5J) revealed that there were ∼0.4 cleavage divisions in the Yki+Stg expression conditions, compared to the ∼3 in controls. During our expansion imaging period, cells in the Yki+Stg animals underwent an average of 3.3 +/-0.97 divisions compared to 3.3 +/-0.80 divisions in wild-type animals (Figures 5K-N). However, we note that in the Yki+Stg animals, expansion started earlier than in the control judging by the temporal offset in SOP specification (Figures S5I, J). In any case, counting the total number of divisions during the cleavage and expansion phases shows that, following larval Yki+Stg expression, the histoblasts undergo an average of 3.7 divisions instead of ∼7 in the controls. Extrapolation of cell number for the anterior nest showed that there were more cells in the Yki+Stg animals than in wild-type, both at the onset of the expansion phase and at the end of tissue expansion, and cells showed a slightly increased apical cell area (Figures 5J and S5M). We also noticed an increase in the number of cells that are extruded from the epithelium upon Yki/Stg overexpression during the expansion phase (Figure S5N). These results show that an increase in the initial cell number and size of the histoblast nests leads to premature proliferation arrest and therefore fewer divisions during the pupal stages.

## DISCUSSION

### Abdominal histoblasts undergo compensatory proliferation

Here, we investigated the cellular mechanisms of compensatory growth by direct observation of this process in the *Drosophila* abdominal epidermis. Whether this tissue is capable of compensatory growth at all has been unclear. Early studies using cauterisation to kill histoblasts during larval development concluded to a weak/absent^35^ or “extensive” ^18^ regenerative capacity based on the recovery of adult cuticle in partially cauterised segments. In contrast, a study using genetics to reduce the number of histoblasts specified during embryogenesis found no evidence for regulation of cell number^36^. The authors used flies expressing 7 copies of the Bicoid (Bcd) morphogen, which reduced the number of histoblasts by ∼38%. Rather than counting the histoblasts, the main assay used to measure the impact on final abdomen cell number was counting the sensory bristles on the dorsal cuticle, which was indeed reduced in the 7xBcd animals^36^. Our data suggest that, in animals where histoblast nest size is reduced prior to expansion, SOP selection is initiated at a time (∼16-20h APF^38^) when cell number is reduced, leading to fewer SOPs being selected and therefore a sparser sensory bristle pattern in adults. In agreement with this, SOP selection occurs in a very reproducible temporal window (^38^; Figure S1) and reducing histoblast nest size by TOR inhibition during larval development leads to sparse sensory bristle distribution in the adult segments (Figure S2E, F). This suggests that counting SOPs is not a reliable proxy for histoblast number. Thus, we conclude that histoblasts are capable of partially regulating their proliferation capacity in response to cell number, allowing for correction of alterations in initial tissue size.

### Cellular mechanisms of compensation

At present, there is limited understanding of the changes in growth kinetics that drive compensatory growth during development. For instance, thymidine incorporation in mouse embryos of reduced size (generated by removing one or two cells at the 4-cell stage) have suggested that compensation can happen both via cell cycle acceleration and divisions beyond the normal proliferative window^9^. Here, we have been able to extract precise growth compensation parameters in live animals. Upon reduction of histoblast nest size, either by preventing the histoblasts from growing during larval development (Figures 2 and 3), or by partially ablating the nests at the onset of metamorphosis (Figure 4), we observe several compensation mechanisms. Most strikingly, in both scenarios the histoblasts can extend the length of their proliferation window by delaying growth termination by several hours. For example, when we ablate all but one cell in the anterior dorsal nest (Figure 4), this cell and its progeny can undergo over 10 rounds of cell division during pupal development, compared with ∼7 for control animals. If we force the histoblasts to divide during the larval stages when they are normally quiescent by expressing Yki+Stg, we can reduce the number of pupal divisions from ∼7 to ∼3.7 (Figure 5). In contrast, we did not observe any adjustment of cell cycle time during the normal proliferative window in response to perturbation. The histoblasts did not accelerate their divisions upon a starting tissue size reduction (Figures 2I, 4C), nor did they slow the cell cycle down when tissue size was increased (Figure 5H). This suggests that the regulatory mechanisms that compensate for histoblast nest size act primarily on the timing of cell cycle exit rather than cell cycle time (Figure 6). It is interesting to note that cell cycle time is largely stable during the expansion phase in control animals (^15^ and Figure 2I), suggesting that physical or metabolic constraints may make it difficult for histoblasts to change their growth and division rates during the normal developmental growth window.

**Figure 6.**
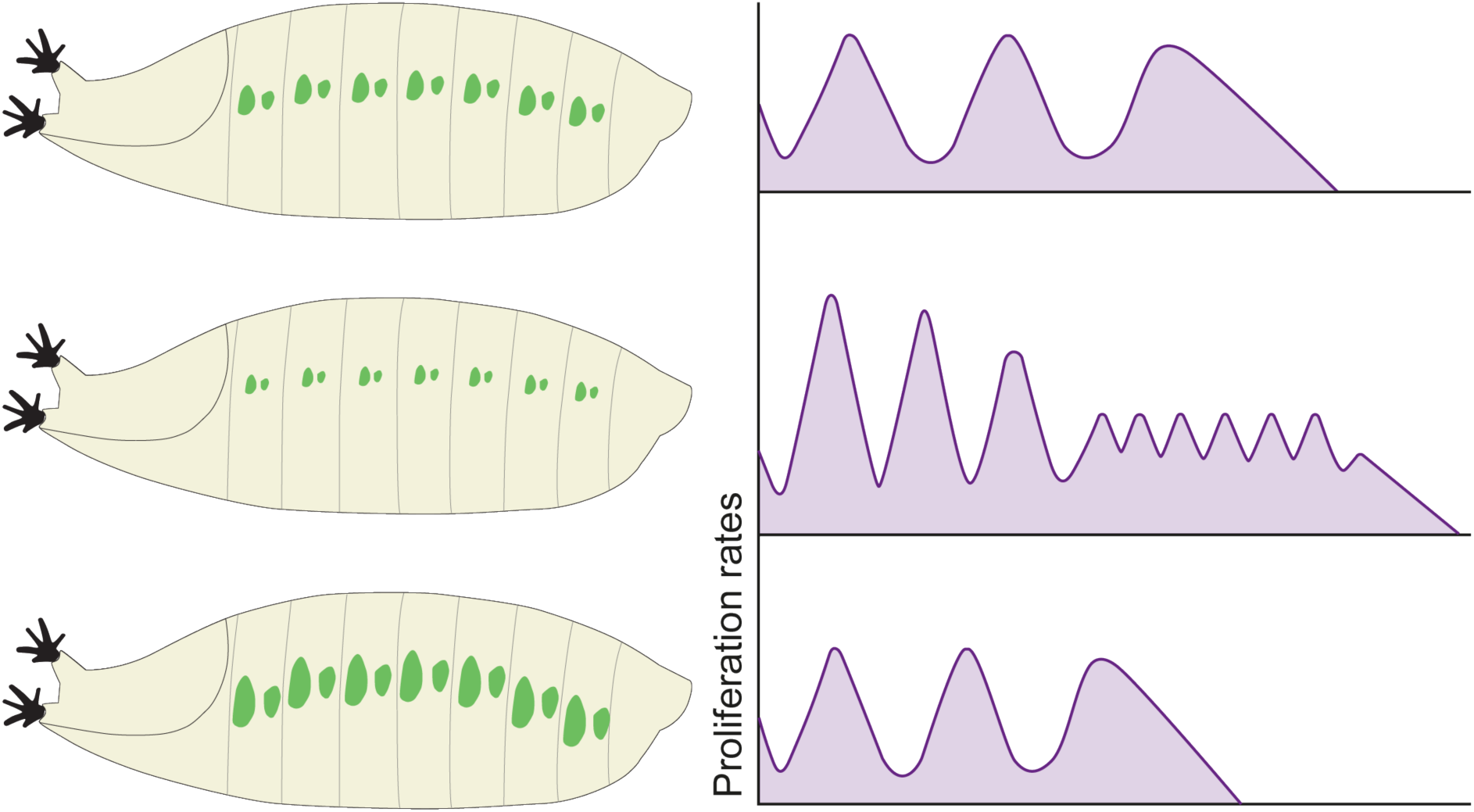
Model for compensatory growth upon changes in tissue occupancy. During normal development, histoblasts proliferate until around 28h APF (top panel). However, if the initial tissue size is reduced, either by reducing cell size or cell number, histoblasts compensate by proliferating for longer to maximise coverage of the abdominal surface (middle panel). On the contrary, if the initial tissue occupies a larger surface, cells stop dividing earlier (bottom panel).

Delaying cell cycle exit is not sufficient to fully compensate for the cell number decrease upon TOR inhibition (Figure 3E) and laser ablation (Figure 4G). In both cases, the histoblasts can increase their apical area to cover a larger surface (Figures 3F and 4H). In addition, we also observed that cells from the ventral histoblast nest appear to be pulled into the dorsal viewing area (Movies S2 and S3). As cell volume is very similar between TOR-inhibited and control histoblasts (Figure 3H), increased apical area upon larval TOR inhibition is not due to a volume increase. Instead, we propose that LEC delamination subjects the reduced histoblast nests to tensile stress, which in turn increases their apical area. Indeed, LEC death has been reported to locally stretch border histoblasts^40,42^. It is less clear why average histoblast surface also increases upon increase in histoblast nest size by Yki+Stg overexpression (Figure S5M). It is possible that the combination of slightly elevated whole pupal volume (Figure S5A), increased extrusion (Figure S5N) and the fact that the abdomen surface appears more convoluted in these animals (Figure 5F, right panel) means that the target surface to be covered by histoblasts is larger than in controls, resulting in a larger apical area.

### Regulatory inputs into histoblast numbers

Can studying compensatory growth inform us about the cues that control growth termination in a wild-type tissue? The ability of histoblasts to change the timing of cell cycle exit depending on starting tissue size suggests that they can sense either tissue volume, or the spatial boundaries of the abdominal segment. This type of tissue size sensing could take the form of a secreted molecule, such as a growth antagonist (chalone) that could stop proliferation once a critical tissue volume is reached^44^. Cell division can be influenced by geometric or mechanical constraints, since cells with a larger surface area or experiencing tensile stress are generally more likely to divide than cells with a smaller area or under compression^37,39,41^. However, this is unlikely to be the spatial signal in the abdominal epidermis, because there is no correlation between histoblast geometry and cell cycle time, and tensile stress does not decrease in this tissue during growth termination^15^. Our laser ablations (Figure S3G) and published cauterisation studies^18,35^ indicate that the histoblasts respect the intersegmental boundaries and the dorsal midline. Indeed, cauterisation of the intersegmental boundaries causes invasion of histoblasts into the neighbouring segment and patterning alterations as anterior and posterior cells from neighbouring segments come into contact^18,35^. The intersegmental LECs, which correspond to the larval muscle attachment sites, are the last LECs to disappear^17,18,35,49^, and present a distinctive chain-like morphology that separates the segments (Figure 1F and Movie 1). It is therefore possible that these boundary cells act as spatial organisers of proliferation, either by preventing premature mixing of histoblasts from neighbouring segments, or by actively signalling to the histoblasts.

Although reduction in starting tissue size leads to a delay in cell cycle exit, the cell cycle progressively slows down as the histoblasts divide beyond their normal proliferative window (Figures 2I and 4C). Thus, as development proceeds, histoblasts become less able to divide, suggesting a temporal input into growth termination. One possible mechanism may be limiting nutrient availability. During pupal life, the animal is not feeding and therefore relies on stores accumulated during larval stages to support metamorphosis^43,52^. It is therefore conceivable that nutrients become scarce as pupal development proceeds, providing a temporal brake on histoblast proliferation. Alternatively, hormonal changes such as increase in the steroid hormone ecdysone could provide temporal inputs into histoblast proliferation^53,54^. In wing imaginal discs, both ecdysone and the nutrient-sensitive TOR pathway have been implicated in cell cycle arrest at the end of larval development^55,56^. However, in histoblasts, reduction of TOR activity during the expansion phase did not appear to affect abdominal segment size (Figure S2), suggesting distinct mechanisms may be at play in this system.

Interestingly, although cell cycle time remains mostly constant during histoblast expansion (^15^ and Figure 2I), we consistently observe a slight increase after 26h APF when the majority of cells have arrested (Figure 2J). Upon TOR inhibition (Figure 2I) or laser ablation (Figure 4C), cell cycle times closely mirror this increase, before further rising as cells proliferate long beyond the normal temporal window. This cell cycle slowdown in controls may therefore represent the temporal regulatory input starting to come into play before another (spatial) signal leads to cell cycle exit, consistent with the possibility that at least two signals are functioning redundantly to promote timely histoblast growth termination. Redundancy in growth termination signals is an attractive model, given the remarkable precision of size control *in vivo*^1,2^, and identifying these signals will be essential to unravel how tissue and body size are determined.

## ACKNOWLEDGEMENTS

We thank Y. Bellaïche, A. Teleman, the Bloomington *Drosophila* Stock Centre, the Vienna *Drosophila* Stock Center and the Kyoto Stock Center for fly stocks. We are grateful to the Crick Advanced Light Microscopy facility and the Crick Fly Facility for support, M. B. Smith for help with the U-Net Neural Network, and J.J. Williamson for help with Tissue Miner. We are grateful to L. Fabrizio, L. Requena, B. Tapon, and Y. Wu for helping with skeleton correction. We thank A. Moraiti and J.P. Vincent for critical reading of the manuscript. AF was funded by the European Union’s Horizon 2020 research and innovation programme under the Marie Skłodowska-Curie grant agreement MSCA-IF-EF-ST No 795060. This work was supported by a Wellcome Trust Investigator award (107885/Z/15/Z) to NT. Work in the NT and GS labs was supported by the Francis Crick Institute, which receives its core funding from Cancer Research UK (CC2138, FC001317), the UK Medical Research Council (CC2138, FC001317), and the Wellcome Trust (CC2138, FC001317). For the purpose of Open Access, the authors have applied a CC BY public copyright licence to any Author Accepted Manuscript version arising from this submission.

## AUTHOR CONTRIBUTIONS

Conceptualization: AF, GS, NT

Funding acquisition: AF, GS, NT

Experiments: AF, FM, MVH, AA

Experimental methodology: AF, AA, FM, MVH, NT

Data analysis: AF, BA, AC, FM, MVH, GS, NT

Supervision: GS, NT

Writing – original draft: AF, NT

Writing – review & editing: All authors

## DECLARATION OF INTEREST

The authors declare no competing interests.

## STAR METHODS

### Key Resources Table

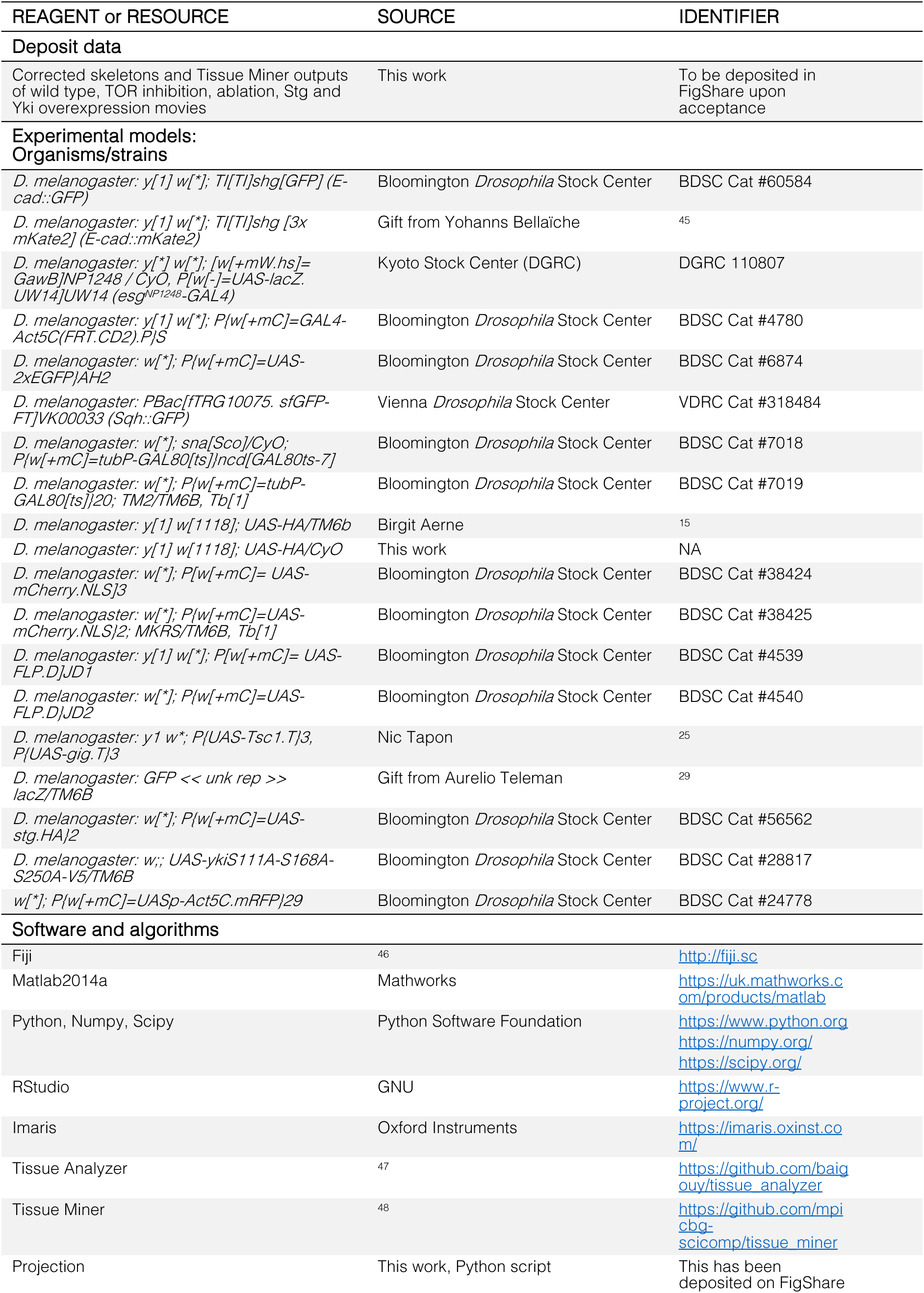
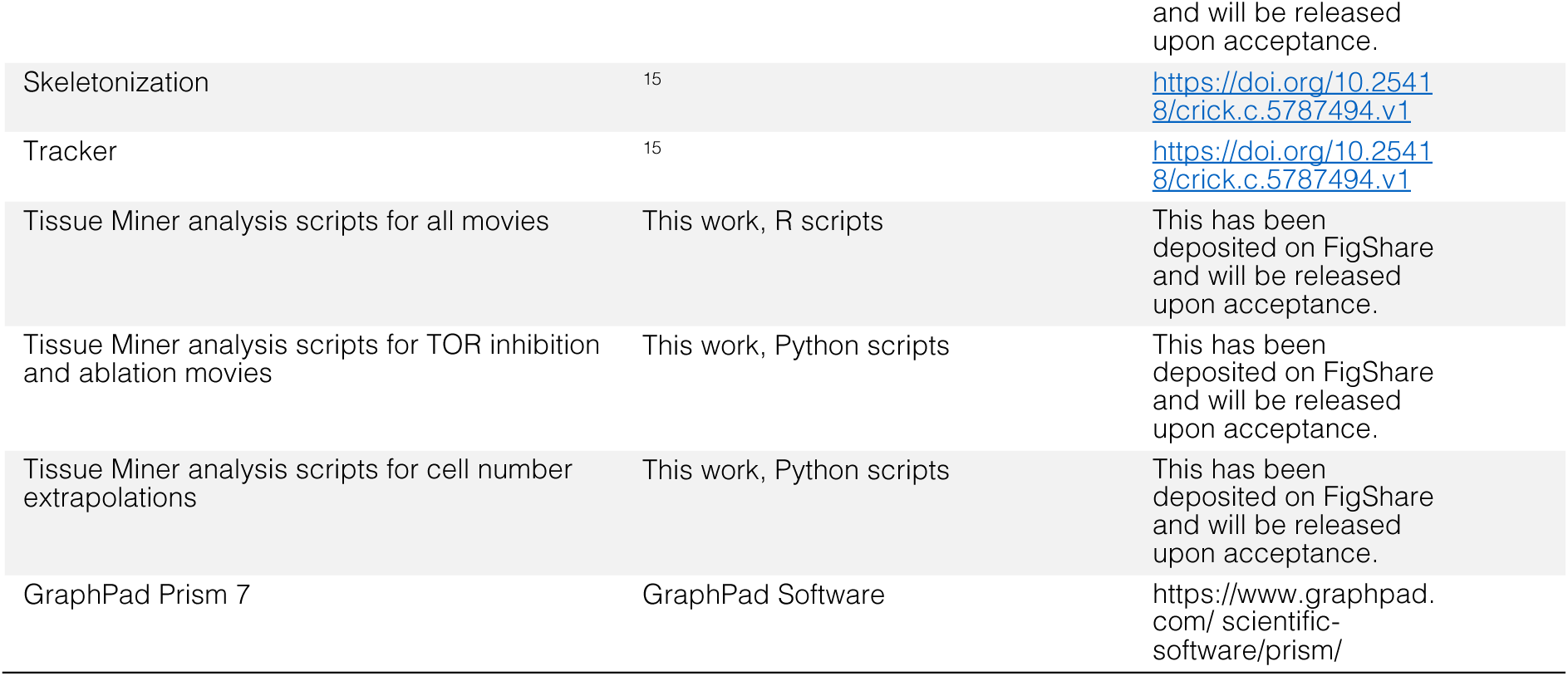

## RESOURCE AVAILABILITY

### Lead contact

Requests for resources, reagents, and further information should be directed to the Nic Tapon.

### Materials availability

All fly stocks are available upon request or from the different *Drosophila* stock centres (see key resources table).

### Data and Code availability

Data and code are available upon request. The scripts for analysis of all experiments are provided free of charge via an online repository (see key resources table).

## EXPERIMENTAL MODEL AND SUBJECT DETAILS

All experiments were performed in *Drosophila melanogaster* (see key resources table and figure genotypes table for details of strains used).

## METHOD DETAILS

### Fly husbandry

Flies were raised at 25°C, unless stated otherwise, on food generated by the Francis Crick Institute Media Facility (33 g agar, 180 g wheat flour, 300 g glucose, 420 g yeast extract, 27 mL propionic and 117 mL nipagin-bavistin mix and 6 L water).

For early prepupal image acquisition, late wandering L3 (wL3) larvae were monitored until pupariation, and white prepupae (0h APF) were collected with a paintbrush. For image acquisition of later pupae, animals were collected before head eversion, kept at 25°C, and then monitored hourly for head eversion to calculate pupal age as 12h APF, four hours prior to imaging at 16h APF. For histoblasts ablation experiments, crosses were allowed to lay eggs at 25°C and then raised at 25°C until 0h APF when animals were subject to laser ablation. Pupae were then allowed to recover at 18°C for around 24h until the start of recording at the age of 16h APF at 25°C.

#### Temporally controlled inhibition of TORC1

We drove the expression of HA (control) or TSC1/TSC2 in the histoblast nests and temporally modulated transgene expression using the GAL4/UAS-GAL80^ts^ or TARGET (temporal and regional gene expression targeting) system^32^. Flies were allowed to lay eggs for 24h at 25°C and then shifted to 29°C for 24h to allow for the *act>CD2>Gal4* cassette to be excised and to ensure a rapid expression of the transgene later in development. Animals were then maintained at 18°C until the time they were shifted to 29°C to induce TOR inhibition until adult eclosion. For the experiment involving TOR inhibition only during the expansion phase, animals were monitored hourly for head eversion to calculate pupal age as 12h APF, four hours prior to imaging at 16h APF at 29°C. For TOR inhibition only during the larval stages, animals were shifted to 29°C after the 24h egg-laying period and shifted to 18°C at the late wL3 stage.

#### Temporal expression of Stg and Yki^3SA^

We drove the expression of HA (control), Stg and/or Yki^3SA^ in the histoblast nests and temporally modulated transgene expression using the GAL4/UAS-GAL80^ts^ or TARGET (temporal and regional gene expression targeting) system^32^. Crosses were kept at 25°C for 24h of egg laying. Larvae were raised at 29°C to induce transgene expression and transferred to 18°C at the wL3 stage to switch off transgene induction. Animals were then monitored hourly for head eversion to calculate pupal age as 12h APF, four hours prior to imaging at 16h APF at 18°C. For early prepupal image acquisition, late wL3 larvae were monitored until pupariation and white prepupae (0h APF) were collected with a paintbrush and imaged at 18°C.

### Imaging

Female pupae were dissected and mounted as described^50^ and the dorsal histoblast nests from the 3^rd^ or 4^th^ segment were imaged. Movies were acquired on a Zeiss LSM 880 confocal microscope at 25°C with a Plan-Apochromat 40x/1.3 oil DIC M27 objective, unless otherwise stated. Images were acquired as two tiles of 1024×1024 pixels with a 10% overlap and the tiled stacks were fused in Zen Blue software. Wild-type, TOR inhibition and nest ablations movies with labelled *E-cad::GFP* were acquired at 25°C with a frame every 2.5 or 5 minutes with 20-30 z-slices, 1μm apart. Volume movies (*EGFP, E-cad::mKate2*) were acquired at 25°C with a frame every 20 minutes with 60-90 z-slices 0.49μm apart. TOR inhibition during the expansion phase movie was acquired at 29°C with a frame every 5 minutes with 20-30 z-slices 1μm apart. Wild-type and TOR inhibition *unk-GFP*, *E-cad::mKate2* movies were acquired at 25°C with a frame every 20 minutes with 30-45 z-slices 1μm apart. Stg, Yki^3SA^, Yki^3SA^+Stg, and HA (control) movies were acquired at 18°C with a frame every 5 minutes with 20-30 z-slices 1μm apart. Wild-type and TOR inhibition early pupal movies were acquired at 25°C with a frame every 10 minutes with 85-110 z-slices with 0.49μm apart. HA or TOR inhibition movies with *E-cad::mKate2, sqh::GFP* were acquired at 25°C with a frame every 20 minutes with 60-100 z-slices 0.50μm apart. Brightfield adult and pupal pictures were acquired on a Zeiss Discovery V20 stereo microscope.

### Laser ablations

Movies were acquired at 25°C on a Zeiss LSM 780 confocal microscope with a Coherent Chameleon NIR tunable laser. For histoblast ablation a Plan-Apochromat 40x/1.3 oil DIC M27 objective was used. Cells were ablated with a wavelength of 780nm at 35% power with the ablation taking 0.254ms.

### Movie segmentation and analysis

Image processing was performed on projected cell surfaces which were subsequently segmented (skeletonized) before being tracked. All these steps were done using a previously generated pipeline developed in the laboratory^15^ with a few modifications outlined in Methods S1. Tissue Miner was then used to extract data from tracked cells^48,51^, as done previously for wild-type movies^15^.

### Tissue and cell volume measurements for growth rate quantification

For analysis during the prepupal stage, we drove the expression of *UAS-mCherry.NLS* to count the number of cells in the anterior and posterior dorsal nests and *UAS-EGFP* was co-expressed to allow tissue volume quantification. To count cell numbers for each timepoint, we used the Cell Counter ImageJ/Fiji Plugin (K. de Vos). For volume quantification of the histoblasts during the expansion phase (16h - 36h APF) we drove the expression of *UAS-EGFP* throughout development while imaging cell junctions with *E-cad::mKate2*. We used the cell junction marker to process and analyse the movies with Tissue Miner^48,51^. The projected cell surfaces were segmented, manually corrected, and further tracked, allowing subsequent analysis.

Using the skeletonized image, we drew the outline of the entire tissue and of the ‘noborder Region of Interest (ROI)’ to specifically quantify growth rates during this developmental stage. This ROI only considers those cells we can follow during the entire movie and using the outline of the ROI, we removed the fluorescence signal from those cells that either leave the field of view later in the movie or that come into the field of view from the neighbouring segments. We overlaid the skeletonised image with the EGFP image to delete what is not part of the ROI. This was done for each timepoint separately using a custom-made ImageJ/Fiji macro.

In both early prepupal movies as well as later pupal movies, EGFP images were binarized and saved as individual timepoints. These images were then imported to Imaris 9.5.1 for surface processing (Figure S1A; Video S1). In Imaris, the Surface tool was used to quantify tissue volume. The surface detail value used was 0.415μm with absolute intensity used for thresholding. The lower limit threshold value was then manually adjusted. After generating the final surface, a step of manual processing was carried out (using the Edit tab) so that non-histoblast tissue expressing EGFP was excluded from the analysis (such as removing the underlying hemocytes). Using the Statistics tab in Imaris, we extracted the tissue volume values for each timepoint. We then calculated the average cell volume by dividing the total volume by the number of cells for each timepoint. Relative growth rate was calculated as the tissue/cell volume difference between two timepoints divided by the tissue/cell volume at the first time point, divided by the time difference between the two timepoints.

### sqh::GFP movies to extrapolate cell volume using tissue height

The *E-cad::mKate2* cell junction marker was used to quantify cell number, tissue area, and cell area. The projected cell surfaces were segmented, manually corrected, and further tracked allowing subsequent analysis with Tissue Miner^48,51^. We used the *sqh::GFP* images to quantify tissue height every hour. For each timepoint, we used six cross-sections along the dorso-ventral axis to measure the height of the histoblast nests. For consistency, the positions along the anterior-posterior axis of all cross-sections were the same, starting at y=300 until y=800, with 100 pixels distance between each one (Figure S3J). For each cross-section, we measured the area and the width of the tissue. By dividing the area by the width, we calculate the average height for each cross-section. We obtained the height for each developmental time by averaging the height of the six cross-sections for that specific timepoint. We then calculated the average cell volume by multiplying the height by the average area for each timepoint.

### Quantification of pupal volume

Pictures were taken with a Zeiss Discovery V20 stereo microscope. Measurements were done using ImageJ/Fiji software. The pupal shape was approximated as an ellipsoid of revolution of long axis L and short axis l, and volume was given by the formula 4/3π(L/2)(l/2)^2^. For pupal size measurements, female wL3 larvae were collected and raised separately until pupa formation and volume was then determined as described above.

## QUANTIFICATION AND STATISTICAL ANALYSIS

All statistical tests were performed using R or Graphpad Prism. Statistical tests used and number of repeats are indicated in the figure legends or in the text.

## SUPPLEMENTAL INFORMATION

Document S1. Figures S1-S5.

Methods S1. Supporting information and methodologies. Figure genotype table and movie segmentation and analysis workflow.

Video S1. Tissue volume reconstruction, related to Figures 1 and S1 Video S2. TOR inhibition movies, related to Figures 2 and S2

Video S3. Ablation movie, related to Figures 3 and S4

Video S4. Yki+Stg overexpression movie, related to Figures 4 and S5

---

**Figure S1.**
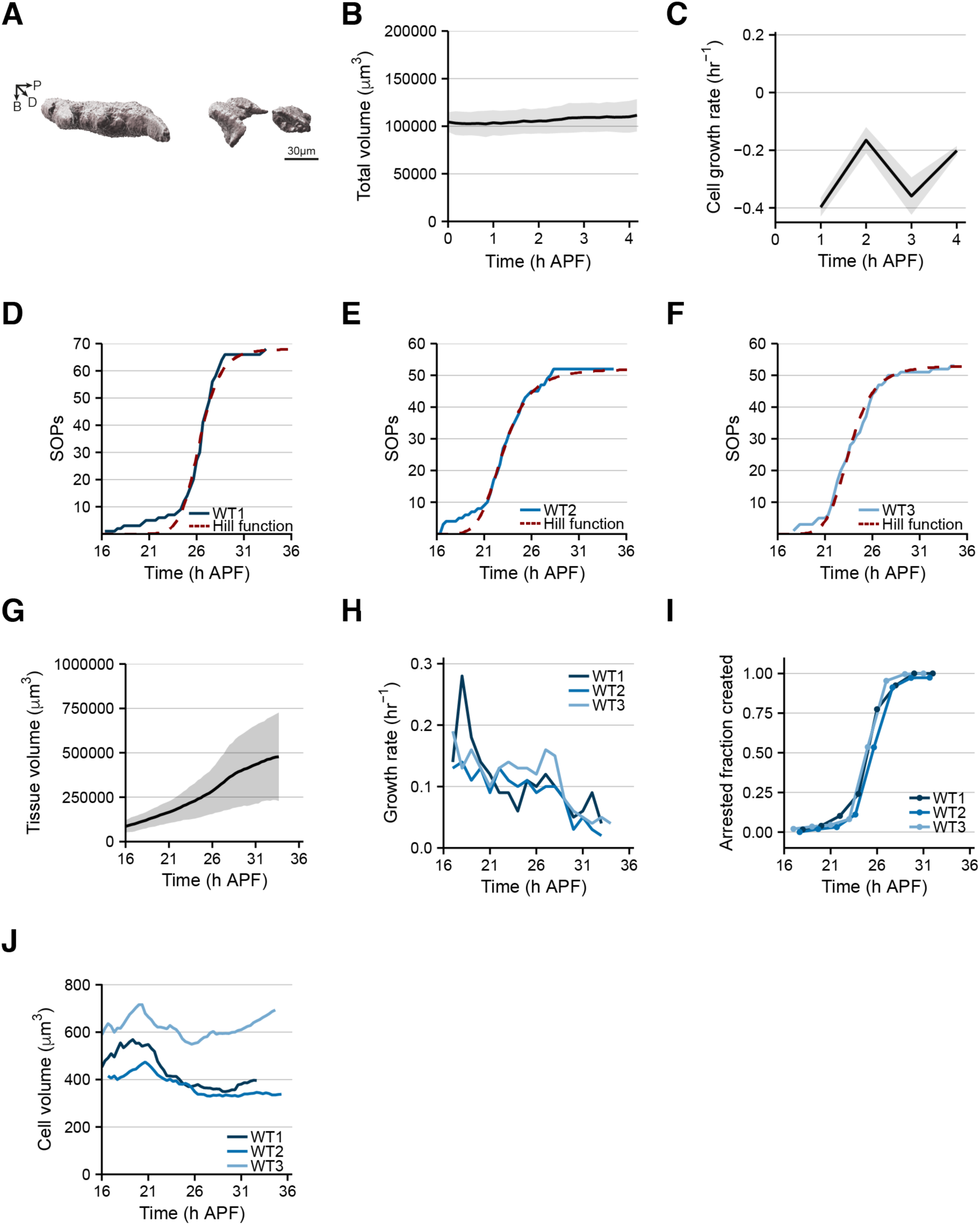
Analysis of histoblast growth dynamics. (A) 3D reconstruction of the histoblast nests at 0h APF. Scale bar = 30 μm. Posterior is to the right, apical to the top and dorsal forward. See also Video S1 and STAR Methods. (B) Total tissue volume as a function of time in wild-type movies at the prepupal stage (n=3). (C) Cell growth rate as a function of time (n=3) during the prepupal stage. Cell growth rate was calculated as the cell size difference between two timepoints divided by the initial cell size. (D-F) Time alignment method for multiple movies. Normalized count of the appearance of sensory organ precursors (SOPs) over time. Each movie is time-aligned relative to movie WT1 (whose first frame is set to be at 16h APF), by an offset determined by fitting Hill functions to the individual movie data shown in (D-F). See also Methods S1. (G) Average total tissue volume as a function of time of the expansion phase in wild-type movies (n=3). Data from noborder ROIs. (H) Growth rate as a function of time for the three wild-type movies analysed. Growth rate was calculated as noborder ROI tissue size difference between two timepoints divided by initial tissue size. Data from noborder ROIs. See also Figure 1, Video S1 and STAR Methods. (I) Fraction of arrested cells as a function of time, for the three wild-type movies analysed. Data from noborder ROIs. (J) Cell volume as a function of time for the three wild-type movies analysed. Cell volume was calculated by dividing the total volume by the number of cells for each timepoint. Shaded grey error bars represent SD.

**Figure S2.**
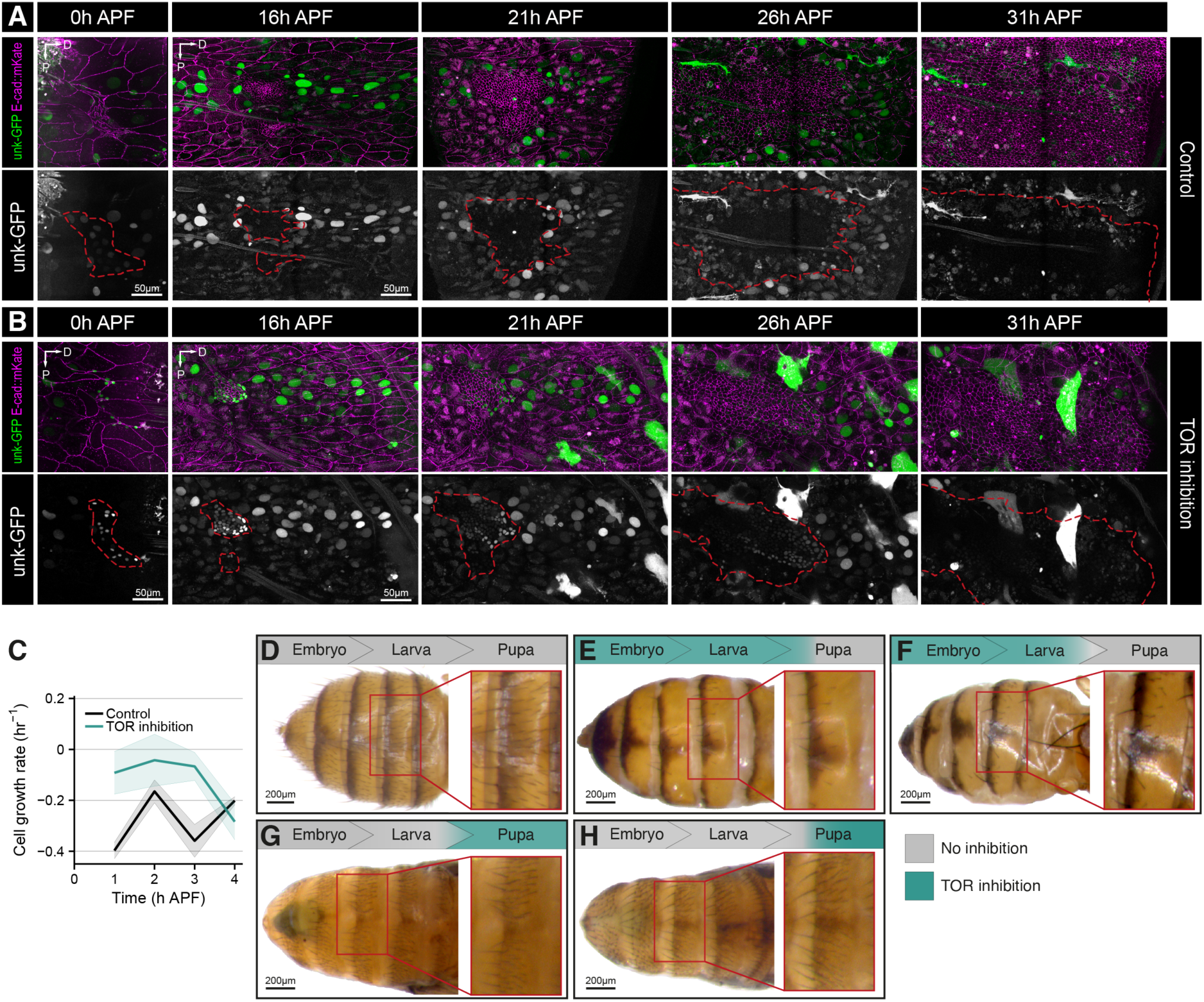
TOR is only required during the larval stages for histoblast growth. (A and B) Confocal images of histoblasts expressing *E-cad::mKate* (in magenta on the top panels) and the *unk-GFP* reporter (in green on top panels and white on the bottom panels) at the time points indicated in control animals (A) or upon inhibition of TOR (B). Histoblasts are outlined in red. Scale bar = 50 μm. Dorsal is to the right; anterior is to the top. (C) Average cell growth rate as a function of time during the prepupal stage for wild-type and TOR inhibition movies (n=3). Cell growth rate was calculated as cell size difference between two timepoints divided by initial cell size. Error bars represent SD. (D-H) Dorsal view of adult (D-F) or pharate adults (G, H) abdominal cuticles of wild-type animals (D), upon inhibition of TOR signalling (E-H) from the embryonic stage until ∼20h APF (E), from the embryonic stage until the end of the larval stages (F), only during the pupal stage (G) or only during the expansion phase of histoblast development (H). The left panels show the female abdomens, and the right panels show an inset at higher magnification of a smaller region of the abdomen. For each panel, a schematic depicting the time of TOR inhibition (in green) or no inhibition (in grey) is shown above. In all cases, the adult cuticle covers the dorsal surface, suggesting the histoblast coverage was largely normal. Note however the sparse implantation of bristles in E and F. Animals shown in G and H are pharate adults that did not hatch, likely due to inhibition of TOR in other tissues than the abdomen. Scale bar = 200 μm. Anterior is to the left. See STAR Methods.

**Figure S3.**
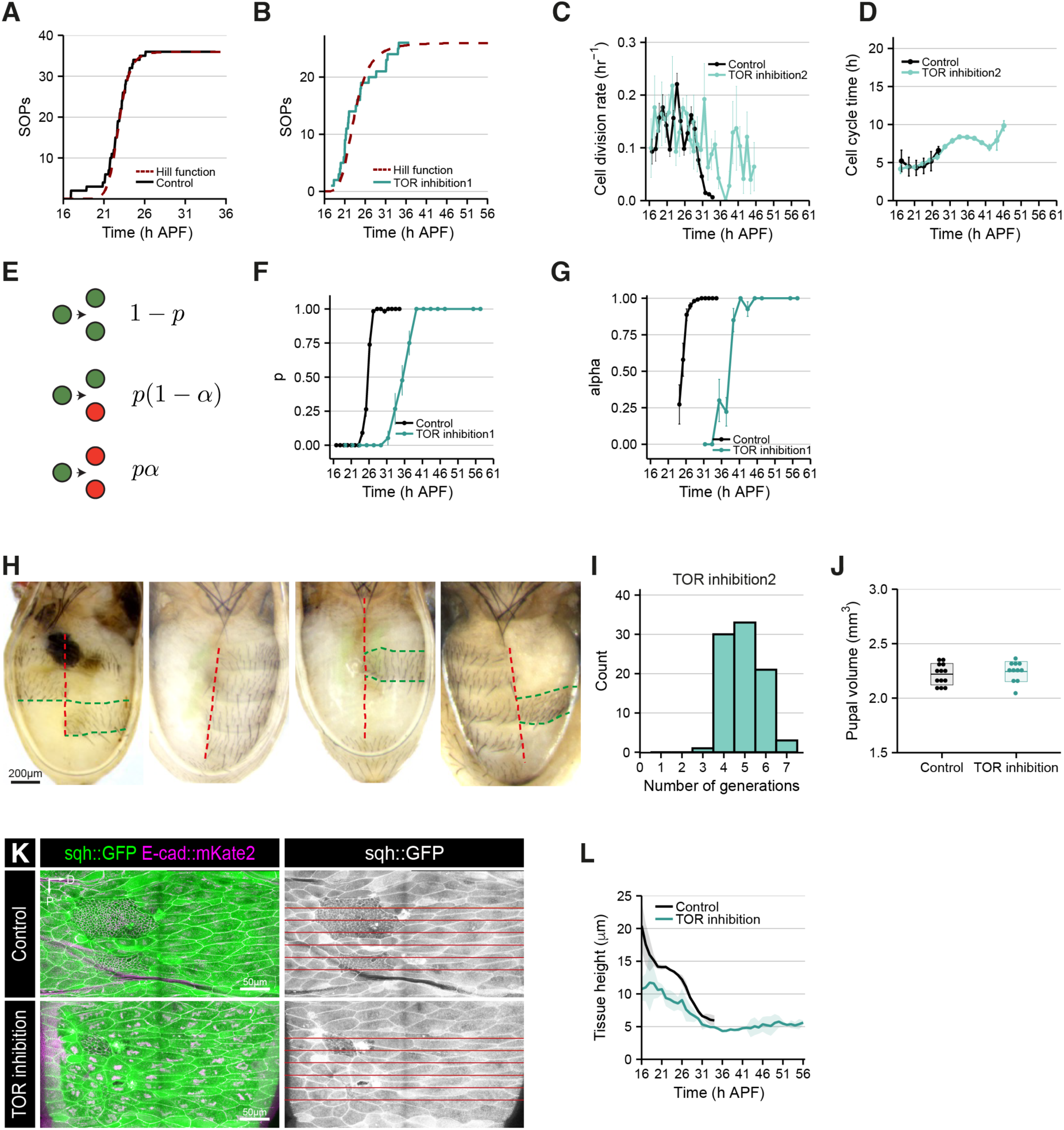
Effects of TOR inhibition on growth dynamics during the expansion phase. (A and B) Time alignment method for wild-type and TOR inhibition movies. Normalized count of the appearance of sensory organ precursors (SOPs) over time in wild-type (A) and TOR inhibition (B) movies. Each movie is time-aligned relative to the wild-type movie (whose first frame is set to be at 16h APF), by an offset determined by fitting Hill functions to the individual movie data shown in (A-B). See also Methods S1. (C) Cell division rate in a wild-type and a TOR inhibition movie as a function of time. Data from noborder ROIs. See also Figure 2 and Methods S1. This movie (TOR inhibition 2) was shorter than the movie presented in Figure 2 (TOR inhibition 1), therefore we could not calculate the fraction of arrested cells, as the cell division rate had not reached 0 by the end. Nevertheless, the movie recapitulates all key aspects of the TOR inhibition phenotype: prolonged proliferation window (Figure S3C), increased cell cycle time at later time points (Figure S3D), and increased cell area (Figure 3F) were confirmed in this independent movie. (D) Cell cycle time as a function of developmental time for wild-type and TOR inhibition movies. The dots represent the binned data, and the error bars are the SEM for the bin. Data from noborder ROIs. (E) Schematic defining the parameters ρ and α characterizing arrested cell creation. ρ is the probability that a division creates at least one arrested cell and α is the probability that a division creates two arrested cells, conditioned on the cell division giving rise to at least one arrested cell. (F) Probability ρ that a division creates at least one arrested cell as a function of time for wild-type and TOR inhibition movies. (G) Probability α that a cell division gives rise to two arrested cells, conditioned on the cell division giving rise to at least one arrested cell for wild-type and TOR inhibition movies. (H) Dorsal view of adult abdominal cuticles upon ablation of histoblast nests at 0h APF from different abdominal segments. In all the cases, the remaining histoblasts do not cross over the intersegmental or dorsal midline boundaries. Red and green dashed lines indicate the dorsal midline and the intersegmental boundaries, respectively. Scale bar = 200 μm. (I) Histogram showing the distribution of histoblast generation numbers during the expansion phase for a TOR inhibition movie. (J) Pupal volume upon overexpression of HA (control) or TSC1/TSC2 (TOR inhibition) in the histoblast nests (*esg^NP^*^1248^–*Gal4* driver). (K) Example confocal images of histoblasts expressing *sqh::GFP* (in green on left panels and in grey on right panels) and *E-cad::mKate* (in magenta on left panels) in control animals (top panel) or in animals expressing TSC1/TSC2 in the histoblasts (bottom panel). The red horizontal lines indicate the position of the cross-sections we used to measure tissue height for each developmental timepoint. Scale bar = 50 μm. Dorsal is to the right; anterior is to the top. (L) Average tissue height as a function of time for control and TOR inhibition movies (n=2). Error bars represent SD.

**Figure S4.**
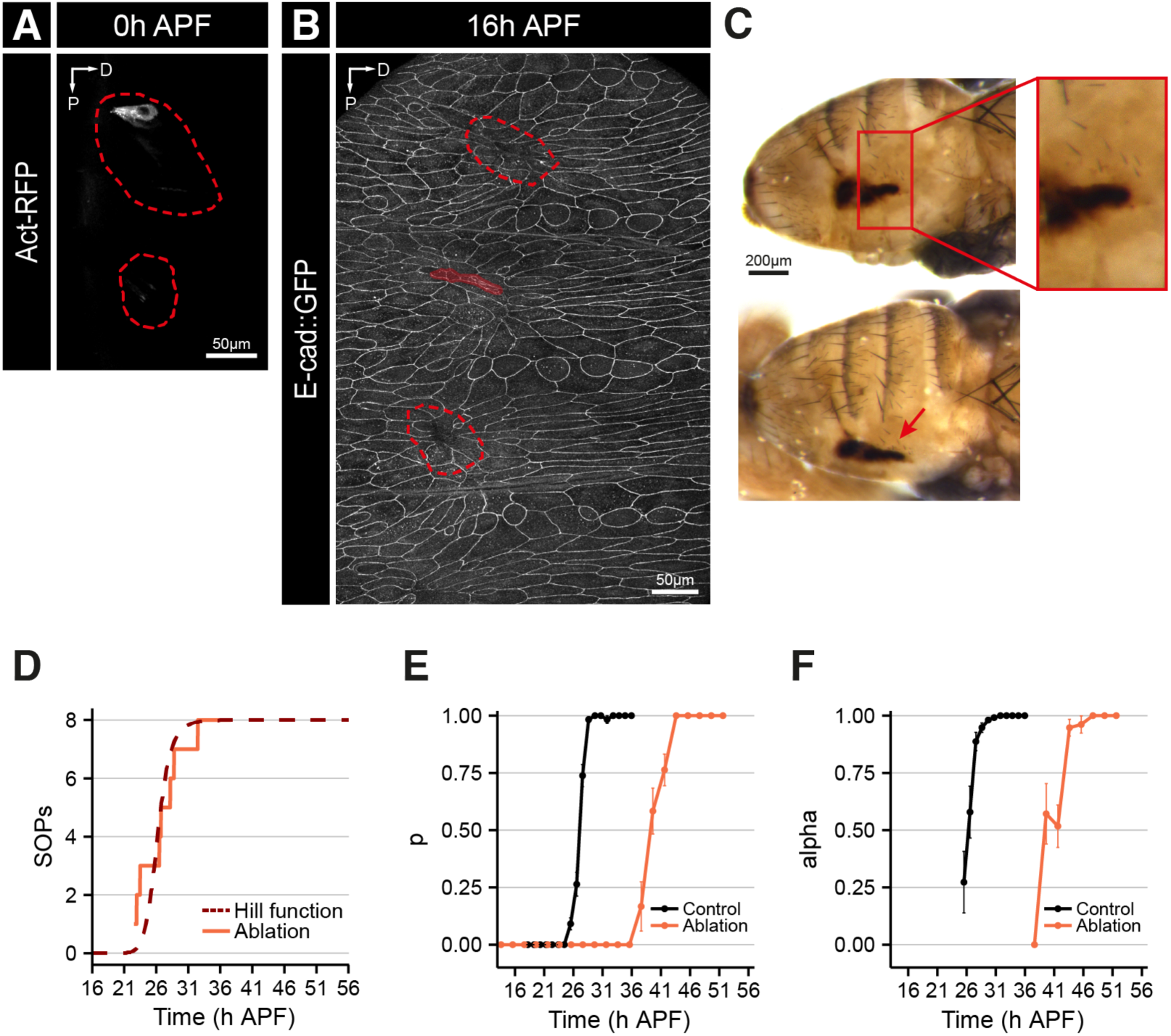
Ablation of most anterior nest cells. (A) Confocal image of histoblast dorsal nests at 0h APF upon ablation of all cells from both anterior and posterior nests except for one cell left in the anterior nest. Histoblasts are labelled by driving *Act-RFP* (in grey). Dashed red line indicates where the nests would be. Scale bar = 50 μm. Dorsal is to the right; anterior is to the top. (B) Confocal image of histoblast dorsal nests expressing *E-cad::GFP* at 16h APF upon ablation of all cells except for one anterior cell in the 4^th^ segment. All histoblasts of neighbouring segments were also ablated. Scale bar = 50 μm. (C) Dorsolateral (top panel) and dorsal (bottom panel) views of the resulting adult abdominal cuticle after ablation as done in A and B. Right panel shows an inset at higher magnification of a small region of the abdomen showing that although a scar due to the laser injury remains, adult cuticle bearing sensory bristles has been secreted by the remaining histoblasts. Scale bar = 200 μm. (D) Time alignment method for the ablation movie. Normalized count of the appearance of sensory organ precursors (SOPs) over time for the ablation movie. The movie is time-aligned relative to the wild-type movie in Fig S3A (whose first frame is set to be at 16h APF), by an offset determined by fitting Hill functions to the individual movie data shown in (D). See also Methods S1. (E) Probability ρ that a division creates at least one arrested cell as a function of time for wild-type and ablation movies. (F) Probability α that a cell division gives rise to two arrested cells, conditioned on the cell division giving rise to at least one arrested cell for wild-type and ablation movies.

**Figure S5.**
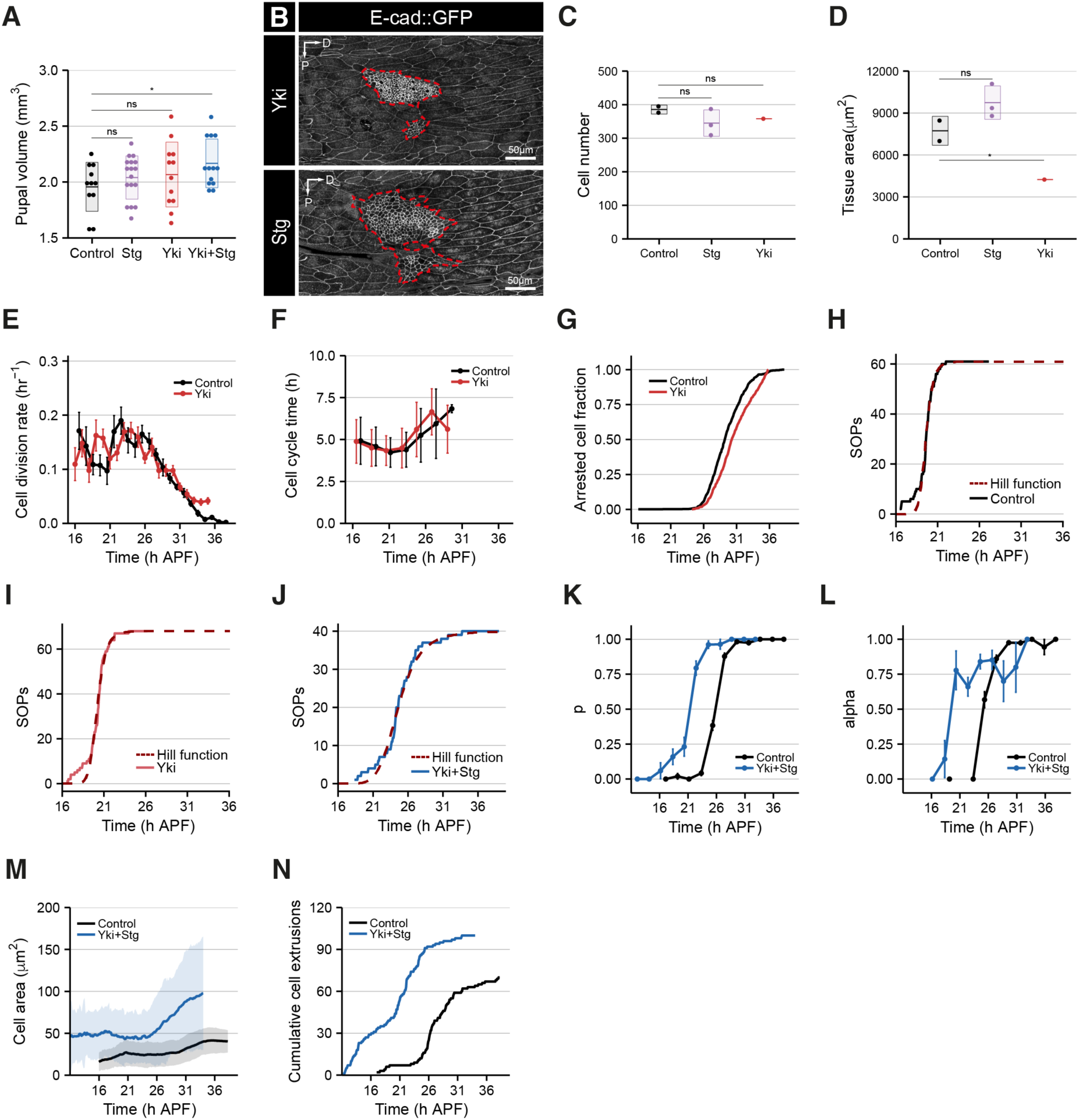
Effect of overexpression of Yki and/or Stg on histoblast growth dynamics. (A) Pupal volume upon overexpression of HA (control), Stg, Yki and Yki+Stg in the histoblast nests (with the *esg^NP^*^1248^–*Gal4* driver and *tub-GAL80^ts^*) from the embryo until the end of the larval stages. (B) Time-lapse confocal images of pupae expressing *E-cad::GFP* at 16h APF upon expression of Yki (top) or Stg (bottom) during the larval stages. Histoblasts are outlined in red. Scale bar = 50 μm. Dorsal is to the right; anterior is to the top. (C) Cell number of dorsal histoblasts nests at 16h APF upon overexpression of HA (control), Stg, and Yki as in (A). (D) Tissue area of dorsal histoblasts nests at 16h APF upon overexpression of HA (control), Stg, and Yki as in (A). (E) Cell division rate in wild-type and Yki overexpression movies as a function of time. Data from noborder ROIs. Error bars represent SEM. (F) Cell cycle time as a function of developmental time for wild-type and Yki overexpression movies. The dots represent the binned data, and the error bars are the SEM for the bin. Data from noborder ROIs. (G) Fraction of arrested cells as a function of developmental time, for wild-type and Yki overexpression movies. Data from noborder ROIs. (H-J) Time alignment method for HA (H), Yki (I) and Yki+Stg (J) overexpression movies. Normalized count of the appearance of sensory organ precursors (SOPs) over time for the HA, Yki and Yki+Stg overexpression movies. The Yki (I) movie is time-aligned relative to the control (HA) movie (whose first frame is set to be at 16h APF), by an offset determined by fitting Hill functions to the individual movies. For the Yki+Stg experiment, the HA (H) movie is time-aligned relative to the Yki+Stg (J) movie (whose first frame is set to be at 11h APF), by an offset determined by fitting Hill functions to the individual movie data shown in (H and J). See also Methods S1. (K) Probability ρ that a division creates at least one arrested cell as a function of time for wild-type and Yki+Stg overexpression movies. (L) Probability α that a cell division gives rise to two arrested cells, conditioned on the cell division giving rise to at least one arrested cell for wild-type and Yki+Stg overexpression movies. (M) Apical cell area as a function of time for both wild-type and Yki+Stg overexpression movies. Data from noborder ROIs. Error bars represent SD. (N) Cumulative number of cell extrusions as a function of time for both wild-type and Yki+Stg overexpression movies.

## Notes

### Competing Interest Statement

The authors have declared no competing interest.

